# Resource and seasonality drive interspecific variability in a Dynamic Energy Budget model

**DOI:** 10.1101/2021.06.02.446572

**Authors:** Joany Mariño, Suzanne C. Dufour, Amy Hurford, Charlotte Récapet

## Abstract

Animals show a vast array of phenotypic traits in time and space. These variation patterns have traditionally been described as ecogeographical rules; for example, the tendency of size and clutch size to increase with latitude (Bergman’s and Lack’s rules, respectively). Despite considerable research into these patterns, the processes behind trait variation remain controversial. Here, we show how food variability, which determines individual energy input and allocation trade-offs, can drive interspecific trait variation. Using a dynamic energy budget (DEB) model, we simulated different food environments as well as interspecific variability in the parameters for energy assimilation, mobilization, and allocation to soma. We found that interspecific variability is greater when the resource is non-limiting in both constant and seasonal environments. Our findings further show that individuals can reach larger biomass and greater reproductive output in a seasonal environment than in a constant environment of equal average resource due to the peaks of food surplus. Our results agree with the classical patterns of interspecific trait variation and provide a mechanistic understanding that supports recent hypotheses which explain them: the resource and the eNPP (net primary production during the growing season) rules. Due to the current alterations to ecosystems and communities, disentangling trait variation is increasingly important to understand and predict biodiversity dynamics under environmental change.

## INTRODUCTION

The variation in life-history traits in animal species across temporal and spatial scales is vast (Healy et al. 2019). Despite considerable research describing these traits’ patterns of occurrence, a systematic understanding of the underlying mechanisms and processes remains elusive (Gaston et al. 2008). Disentangling and quantifying variation in biological traits is necessary to explain and predict biodiversity dynamics under environmental change (Cardilini et al. 2016, Cabral et al. 2017), and it may aid in answering broad questions that range from the invasive potential of species (Capellini et al. 2015) and the evolution of senescence (Jones et al. 2014), to predicting the influence of stressors on species assemblages (Darling et al. 2012). Hence, it is fundamental to understand the underlying mechanisms giving rise to animal trait variation.

Among animal traits, body size exhibits enormous diversity within orders and narrower clades of animals, and it is thought to play a pivotal role in all individual’s ecological and physiological processes (Yom-Tov and Geffen 2011, Kozlowski et al. 2020). Variation of body size in time and space is assumed to be a product of evolution modulated by the biotic and abiotic environment (Mayr 1956, Millien et al. 2006, Yom-Tov and Geffen 2011). Additionally, body size tends to covary with several life-history and morphological traits (Olalla-Tárraga et al. 2019). For example, birds’ body size is thought to be positively correlated with clutch size (Jetz et al. 2008, Olalla-Tárraga et al. 2019). Although there is extensive evidence describing and supporting spatial patterns in body size and reproductive output at different biological scales, the processes that underpin their variation are not fully comprehended (Gaston et al. 2008).

The consistent variation in animal traits across time and space, both within and among species or clades, forms the basis of ‘ecogeographical rules’ (Boyer and Jetz 2010, McNab 2010). Two of the most frequently explored interspecific patterns are the increase of body size in closely related endotherms (and some ectotherms) with latitude (“Bergman’s rule”), and the tendency of clutch size to increase with latitude (“Lack’s rule”). In general, empirical evidence in endotherms supports both patterns (e.g., for Bergman’s rule see Ashton et al. 2000, Cardillo 2002, Ashton 2002, Freckleton et al. 2003, Blackburn and Hawkins 2004, Rodríguez et al. 2006, 2008, Olalla-Tárraga and Rodríguez 2007, Ramirez et al. 2008, Olson et al. 2009, Morales-Castilla et al. 2012a, b, Clauss et al. 2013, Torres-Romero et al. 2016, Romano et al. 2020 and reviews in Meiri and Dayan 2003, Watt et al. 2010, Huston and Wolverton 2011. For Lack’s rule see Moreau 1944, Lack 1947, Ashmole 1963, Cody 1966, Ricklefs 1980, Kulesza 1990, Iverson et al. 1993, Griebeler and Böhning-Gaese 2004, Evans et al. 2005, Jetz et al. 2008, Mesquita et al. 2016, Meiri et al. 2020 and review in Boyer et al. 2010. But see Meiri et al. 2004, Medina et al. 2007, Olalla-Tárraga and Rodríguez 2007, Gohli and Voje 2016 for contradictory or ambiguous results). Bergman suggested that a larger body is an adaptation to colder environments because larger organisms have a lower surface-area ratio than smaller organisms, which allows them to conserve heat more effectively (Salewski and Watt 2017). Lack attributed the larger clutch size in northern species than those in the tropics to a greater food abundance and longer daylight periods during the breeding period (Lack 1947). These hypotheses, however, remain highly controversial, and several alternative mechanisms have been proposed (Stearns 1976, Searcy 1980, Blackburn et al. 1999, Jetz et al. 2008, Pincheira-Donoso 2010, Meiri 2011, Olalla-Tárraga 2011).

Recent hypotheses have explained the variation in body size, reproductive output, and life-history traits based on food availability (McNab 2010, Huston and Wolverton 2011). In the “resource rule”, Bergman’s and Lack’s patterns are determined by the size, abundance, and availability of food (McNab 2010). The “eNPP rule” (or “Geist’s rule”) further explains that the mechanism driving food availability is the global distribution of net primary productivity during the growing season (eNPP) (Geist 1987, Huston and Wolverton 2011). Thus, both hypotheses agree that, in species or groups of closely related species, the largest and with greater reproductive output will occur where food availability (or eNPP) is highest (McNab 2010, Huston and Wolverton 2011).

According to the resource and the eNPP rules, food availability variation should be the main determinant of energy input (McNab 2010, Huston and Wolverton 2011). Thus, we hypothesize that resource availability drives individual energy allocation trade-offs among different life-history components, particularly body size or mass and reproductive output. Consequently, if food availability is the critical factor determining individual energy allocation, then trait variability would be minimal when the resource becomes limiting, both in a constant or seasonal environment. Hence, for species with limited phenotypic plasticity, we expect that individuals in a low food environment will exhibit more similar body mass and reproductive output than in a higher food environment. Further, when the resource is seasonal, the periods of food surplus should allow individuals to reach a large body size and have greater fecundity, in agreement with the eNPP rule (Huston and Wolverton 2011). Therefore, we predict that individuals in a seasonal environment should reach a larger body mass and have a greater reproductive output relative to the same organism in an environment with the same average but constant resource.

Research into patterns of trait variation (particularly over large spatial scales) typically expose the variation using some measure of central tendency in the trait of interest concerning particular environmental predictors (Gaston et al. 2008). These patterns are generally approached at three main levels: intraspecific, interspecific, and assemblage-based (Gaston et al. 2008, Yom-Tov and Geffen 2011). Intraspecific studies focus on explaining patterns in individual species’ traits (e.g., fecundity according to the food supply in *Daphnia pulex*; McCauley et al. 1990). Interspecific (or cross-species) research concerns the differences in the pattern of variation in the trait of interest among species, usually within the same clade or taxon (e.g., global variation in avian clutch size; Jetz et al. 2008). Finally, the assemblage or community approach describes traits patterns in communities across different places (e.g., the latitudinal variation of body size in *Plethodon* salamanders’ assemblages in North America; Olalla-Tárraga et al. 2010). Gaps and biases in the knowledge of trait data among species make studies considering the interspecific approach scarcer than those regarding intraspecific or assemblage variation (Gaston et al. 2008). Additionally, the methodological distinction between interspecific and assemblage patterns is often overlooked or confused (Gaston et al. 2008, Olalla-Tárraga et al. 2010). Hence, there is still uncertainty in the mechanisms structuring interspecific variation, hindering our capacity to forecast cross-species responses.

Understanding the effect of environmental predictors on the interspecific variation of body size is complicated by several confounding factors. For example, the use of phenotypic data may lead to the inability to discriminate between genetic (adaptive) and non-genetic (plasticity) sources of variation (Stillwell 2010). Additionally, the conditions during growth can have a strong effect on body size, for which it would be necessary to control for the birth year (Yom-Tov et al. 2006, 2010, Hersteinsson et al. 2009). A proposed solution to avoid these issues is to conduct common-garden or reciprocal transplant experiments (Yom-Tov and Geffen 2011). However, these experiments are typically not feasible because they involve large samples and individuals’ long-term monitoring (Teplitsky and Millien 2014). For this reason, we adopted a modelling approach that allowed us to investigate interspecific variation in traits by assessing their physiological origin without such confounding effects. To test our predictions, we used the ecophysiological description of the individuals proposed by Dynamic energy budget (DEB) theory (Kooijman 2010). In the DEB model, the parameters represent the physiological processes that, together with the environmental conditions, result in different individual traits. The combination of parameter values in the DEB model defines species, and because it has been parameterized for over 2000 animal species (see AmP 2021 for the complete list), it allowed us to reproduce the natural variation observed in experimental data across species. In this way, we rely on a mechanistic description of the individual metabolism and energy allocation to quantify the environment’s effect on cross-species traits.

We evaluated interspecific variability in genetically-determined physiological characters (represented by the model parameters for assimilation, mobilization and allocation of energy) by carrying out numerical simulations of the DEB model and quantifying the differences in individual traits (i.e., biomass, maturity, and reproduction). We considered both constant and seasonal resource conditions in order to provide a complete description of the effect of food availability. We found that resource determines the expression of interspecific differences in the DEB model in both constant and seasonal environments. Further, individuals in a seasonal environment reach a larger body mass and have greater fecundity compared to the same individuals in an environment with an equal average but constant food. Thus, our simulations with the DEB model agree with the expectations according to the resource and eNPP rules and provide a mechanism supporting these hypotheses.

## MATERIALS AND METHODS

To test our prediction regarding the role of resource, we represented interspecific differences with different sets of parameter values and conducted simulations of the DEB model assuming different constant environments. To test our prediction about resource seasonality, we simulated the same interspecific variation in the DEB model but assuming a periodically fluctuating resource. Thus, we evaluated the same set of parameters, each representing a species, both with constant and seasonal resource availability. We then compared the dynamics of two species to highlight the differences due to the resource regime.

### The standard DEB model

We focus our analysis on the standard DEB model, which is the simplest non-degenerated model implied by DEB theory and the most commonly analyzed DEB model (Lika et al. 2011a). The standard DEB model applies to heterotrophic animals, and it supposes that the biomass of an individual is partitioned into reserve energy and structural volume (Kooijman 2010). Hence, four state variables describe individuals: energy in reserve (*E*, J), structural volume (*V*, cm^3^), cumulative maturity energy (*E_H_*, J), and cumulative reproduction energy (*E_R_*, J). We assume that individuals release the reproduction energy continually as gametes, regardless of the environmental conditions, to neglect its potential contribution to the individuals’ biomass. To compensate for the different possible reproduction strategies, we measure reproductive output as cumulative reproductive energy.

The DEB model assumes that energy from food is assimilated into the reserve (Fig. 1) through the assimilation flux 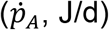. The reserve energy is mobilized according to the mobilization flux 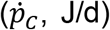. A fixed fraction (*κ*) of the mobilized reserve is allocated to somatic maintenance and volume growth 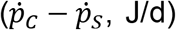. The remaining fraction of the mobilized energy (1 – *κ*) is allocated to maturity maintenance and maturation in juvenile individuals or maturity maintenance and reproduction in adults 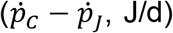. Thus, the temporal dynamic of the individual state variables is:

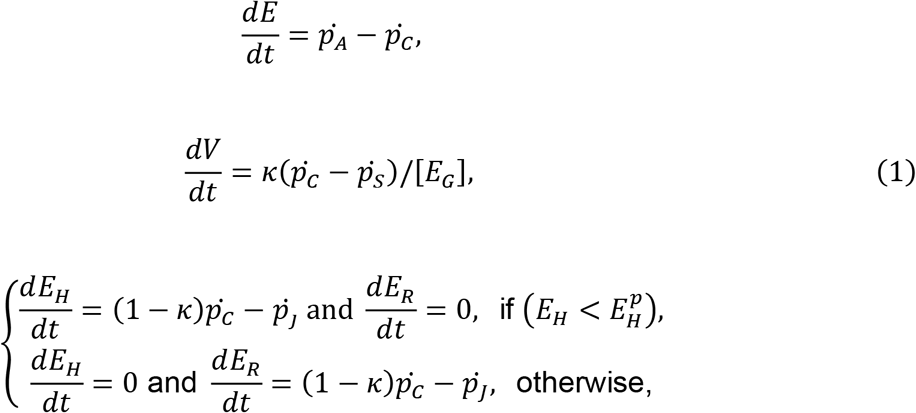

where [*E_G_*] is the volume-specific cost for structure, and the energy fluxes 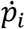 for each process *i* are given in Table 1. The standard DEB model considers organisms with three life stages, as determined by the cumulative maturity level relative to the maturity thresholds parameters, 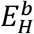 for birth and 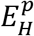 for puberty: embryo (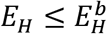 and 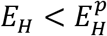), juvenile (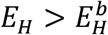 and 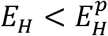), and adult (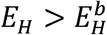 and 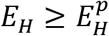).

**Figure 1.**
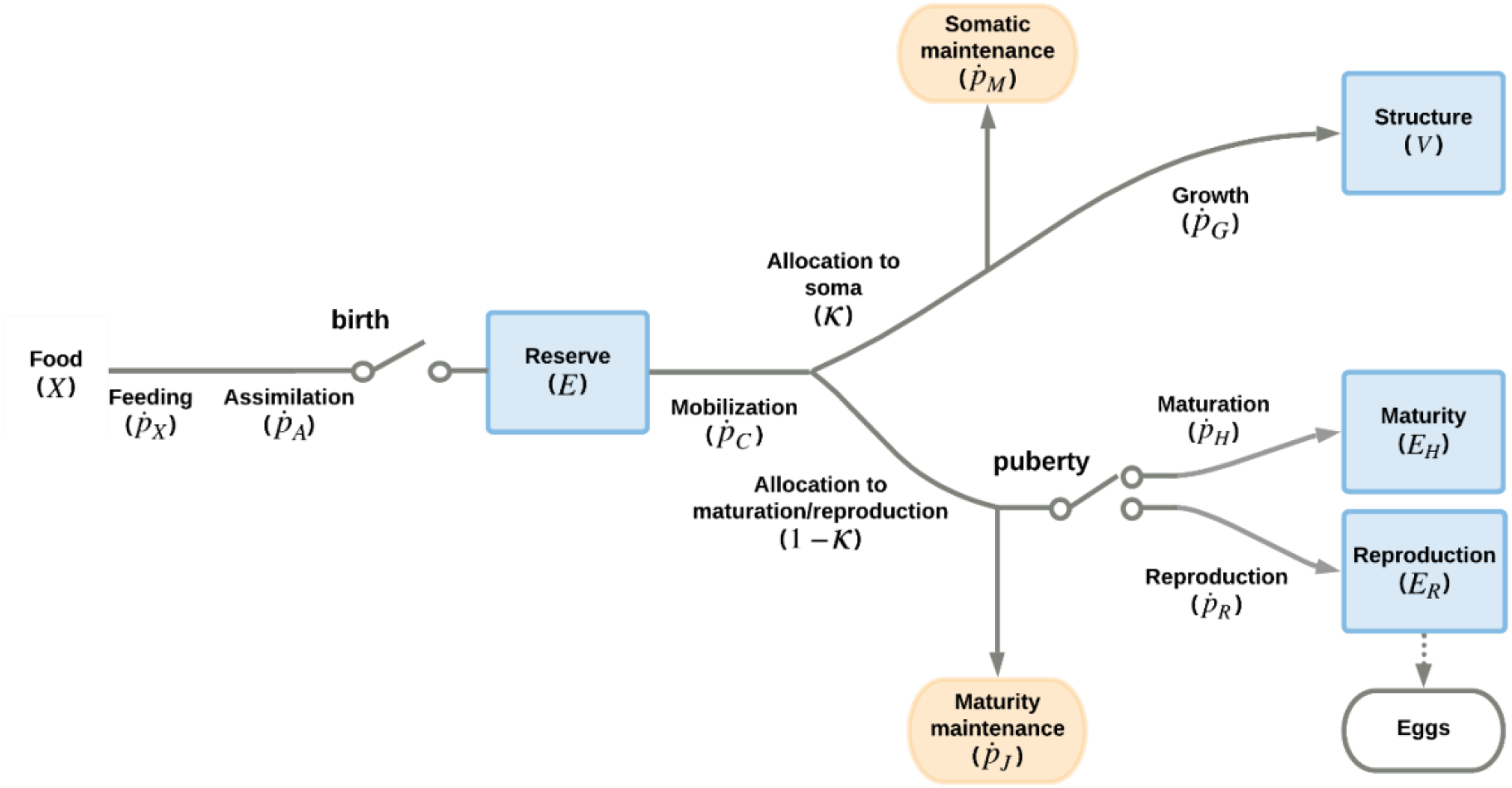
Representation of the standard DEB model (eq. 1). Square boxes denote the state variables, while round boxes represent energy sinks. Lines and arrows correspond to the energy fluxes (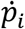 J/d, Table 1). The switches represent metabolic thresholds: birth indicates the start of feeding, while puberty signals the start of energy allocation to reproduction once maturation is complete.

**Table 1.** Energy fluxes (J/d) at each developmental stage. The scaled functional response is *f* (0 ≤ *f* ≤ 1, where 1 is the highest amount of food), 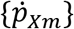 is the maximum surface-area specific ingestion rate 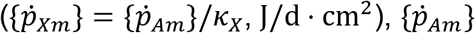 is the maximum surface-area specific assimilation rate (J/cm^3^ · d), *κ_X_* is the assimilation efficiency from food to reserve (dimensionless), 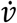 is the energy conductance rate from the energy reserve (cm^3^/d), 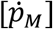 is the volume-specific somatic maintenance cost (J/d · cm^3^), 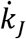 is the maturity maintenance rate coefficient (1/d). Notation: square braces ([]) indicate quantities related to structural volume, curly braces ({}) denote quantities related to structural surface-area, dots (.) indicate rates.

| Flux | Embryo<br>($E_H \leq E_H^b$) | Juvenile<br>($E_H^b > E_H < E_H^p$) | Adult<br>($E_H \geq E_H^p$) |
| --- | --- | --- | --- |
| Feeding, $\dot{p}_X$ | 0 | $f\{\dot{p}_{xm}\}$ | $f\{\dot{p}_{xm}\}$ |
| Assimilation, $\dot{p}_A$ | $\kappa_X \dot{p}_X$ | $\kappa_X \dot{p}_X$ | $\kappa_X \dot{p}_X$ |
| Mobilization, $\dot{p}_C$ | $E \dot{v}(V^{-1/3} - \dot{r})$ | $E \dot{v}(V^{-1/3} - \dot{r})$ | $E \dot{v}(V^{-1/3} - \dot{r})$ |
| Somatic maintenance, $\dot{p}_S$ | $[\dot{p}_M]V$ | $[\dot{p}_M]V$ | $[\dot{p}_M]V$ |
| Maturity maintenance, $\dot{p}_J$ | $\dot{k}_J E_H$ | $\dot{k}_J E_H$ | $\dot{k}_J E_H^p$ |

The mobilization flux 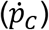 includes the specific growth rate in structural volume:

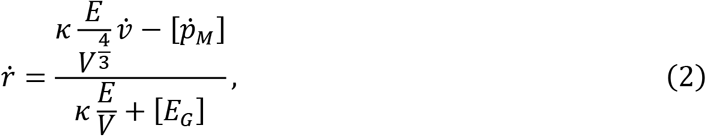

where 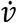 is the energy reserve mobilization rate (cm^3^/d), and 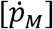 are the volume-specific somatic maintenance costs (J/d · cm^3^).

In the DEB model, both somatic and maturity maintenance have priority over investment in either growth, maturation, or reproduction. Hence, starvation occurs when mobilized energy does not suffice to cover somatic maintenance 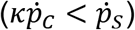 or maturity maintenance 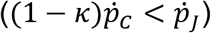. The standard DEB model makes no assumptions about these situations, meaning that organisms follow the same dynamics previously outlined when subjected to starvation. Consequently, when there is prolonged starvation, individuals will degrade the structural mass to cover maintenance costs and shrink in size (because the specific growth rate 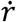 becomes negative).

DEB theory assumes that the metabolic rates are affected by the environmental temperature (Kooijman 2010). Consequently, the model parameters are usually standardized to the reference temperature of 20 °C through the Arrhenius relationship. For simplicity, however, we suppose that the environmental temperature is equal to the reference temperature (*T* = 293.15 K). Further, because food availability often covaries with environmental temperature, we assume that the temperature remains constant and evaluate the effect of seasonality only in the resource.

### Resource

We assumed that the food density operates directly on the scaled functional response *f*, facilitating contrasting the model’s behaviour across different fluctuation regimes (Muller and Nisbet 2000). To investigate the effect of interspecific differences, we first conducted simulations assuming a constant resource, ranging from scarce (*f* = 0.2) to maximum availability (*f* = 1).

To evaluate the effect of resource variability, we assumed a periodically changing functional response, which represents the alternation between two levels of food during the year, similar to a seasonal change. Specifically, the functional response at time *t* oscillates around the average 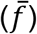 with amplitude *f_a_* and period equal to the length of one year:

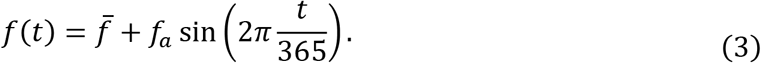

We only consider variations in the value of the mean scaled functional response for simplicity, keeping the amplitude of the oscillations fixed. To assess the initial resource’s effect, we simulated functional responses that start at four different points in the seasonal cycle: maximum resource, intermediate but decreasing resource, minimum resource, and intermediate but increasing resource. These different initial conditions of food can be understood as corresponding to individuals born at different times throughout the year.

### Interspecific variability and parameter space

Despite the diversity of life-history traits and strategies in the animal kingdom, not all strategies are possible (Healy et al. 2019). In order to select a biologically consistent parameter set, we used the AmP collection (a web repository of species parameterized for DEB models; AmP 2021) and the routines in the AmPtool (version 03/2020 AmPtool 2020) MATLAB package (version 9.8, The MathWorks Inc. 2020). We started by taking a subset of the species in the AmP collection (Fig. 2, panel 1). Then we narrowed down the parameter space according to the occurrence frequency of the parameter combinations (Fig. 2, panel 3).

**Figure 2.**
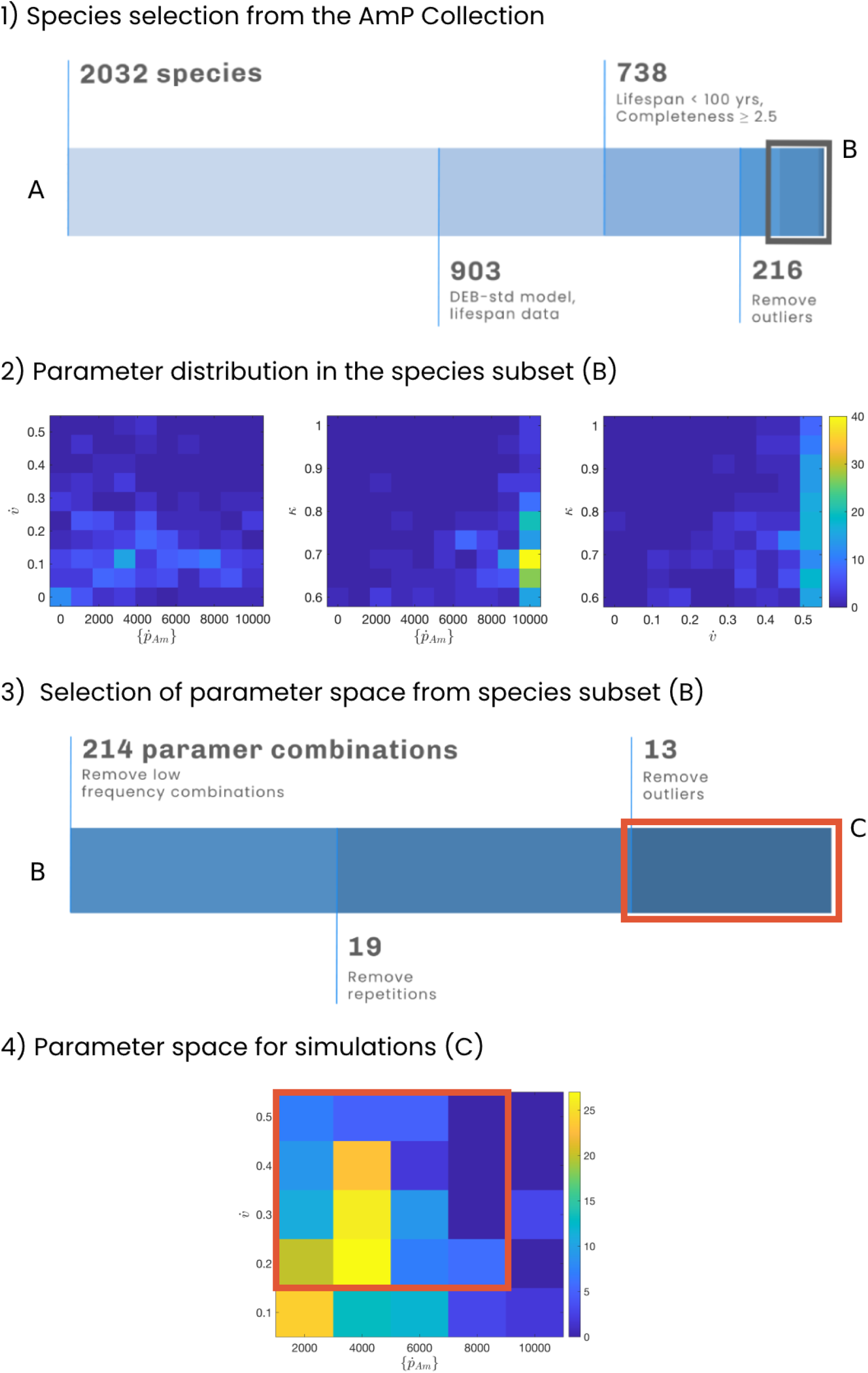
Representation of the selection process that assured biological consistency in the parameter space. Panel 1 shows the species selection in the AmP collection, starting from all the entries (at 03/2020) to a final subset of 216 species (B). The values indicate the number of species at each step. The steps are: i) subset entries modeled with the DEB-std model and containing lifespan data, ii) subset entries with lifespan below 100 years and data completeness equal or greater than 2.5, and iii) remove entries more than 1.5 interquartile ranges above the upper quartile or below the lower quartile of the parameter distribution. In panel 2, the plots show the bivariate distribution of the parameters 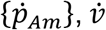, and *κ* in our 216 species subset (B). For all the plots, the parameters’ distribution is not uniform. Hence, all parameter combinations are not equally likely to occur. The colour bar shows the joint frequency of occurrence. Panel 3 shows the selection of the joint parameter space for 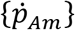 and 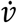 from the species subset B. The values at each step show the number of parameter combinations, starting from 214 to a final subset of 13 parameter combinations (C). The steps are: iv) find the joint distribution of the parameters and remove parameter combinations with a frequency of one, v) remove parameter combinations that are not unique, and vi) remove outlier values. In panel 4, the plot shows the joint distribution of 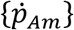 and 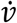 for the 214 parameter combinations more likely to occur in our species subset. The box marks the outer boundary of our parameter space in our parameter combination subset (C). Without repetitions, the parameter space C corresponds to the 16 combinations we evaluated for 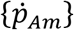 and 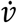.

First, our focus is on the species in the AmP collection (Fig. 2, panel 1). We selected the species modelled using the standard DEB model and containing lifespan data. To assure that the values are consistent, we restricted the entries to those with a lifespan < 100 years and data completeness ≥ 2.5 (the data completeness indicates how much data is available to estimate the DEB parameters, ranging from a minimum of 0 when only maximum body weight or size is known, to a maximum of 10 when all aspects of energetics are known; Lika et al. 2011a). Completeness of 2.5 means that there is data for the species on maximum body weight (or size), age, length and weight at birth and puberty, and growth in time (Lika et al. 2011a); allowing the estimation of the parameters 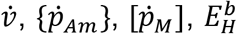, and 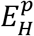 (Kooijman et al. 2008). Then, we removed parameter outliers by excluding values 1.5 interquartile ranges above the upper quartile or below the lower quartile of each parameter distribution, which left 216 species.

Next, we concentrated on the parameters of the 216 species subset. We focused our analysis on three parameters that directly relate to concepts of life-history theory: maximum assimilation rate 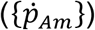, which reflects the ability to acquire energy; energy conductance 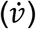, which is related to the “pace-of-life” concept; and energy allocation to maturity and reproduction (*κ*), which reflects the trade-off between somatic growth and reproduction. All combinations of these three parameters in our subset of species are not equally likely to occur, they may not be biologically realistic, and multiple combinations are repeated (Fig. 2, panel 2). For this reason, we calculated the joint distribution of 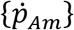 and 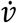 by discretizing their joint range into five intervals and assigning the interval’s mean as the parameter value. Subsequently, we removed the combinations of 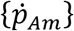 and 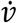 with a low occurrence frequency, leaving the 214 parameter combinations most likely to occur in our species subset (Fig. 2, panel 3). Since these parameter combinations are not unique, we excluded the repetitions, which left 19 unique combinations of 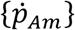 and 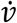. Then, we removed outlier points at the edges of the discrete distribution, leaving a parameter space consisting of 13 combinations of 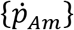 and 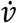. To increase our analysis’s resolution, we added combinations that fell within the parameter space range but had a lower occurrence frequency. Hence, we evaluated 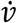 equal to 0.2, 0.3, 0.4, and 0.5, and 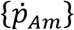 equal to 2000, 4000, 6000, and 8000, which corresponds to 16 different combinations of 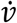 and 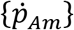 (Fig. 2, panel 4).

Reproduction data is required to estimate the value of *κ*, as well as growth and size at birth and puberty (Lika et al. 2011b). Given that reproduction is often difficult to quantify, many AmP collection entries assume the predefined value of *κ* = 0.8, which results in rapid growth to a large size, long development times, and low reproduction. Even when data is available, due to the simplification of seasonality effects in the data, which do not consider the cycles in up- and down-regulation of metabolism, the parameter estimation is likely to result in a high value for *κ* (Kooijman and Lika 2014). Hence, the AmP collection is biased to high values for *κ*. However, it has been shown that a lower value of *κ* (*κ* ≤ 0.5) is likely to fit growth and reproduction data equally well as the larger value, and producing individuals with reduced growth and reproduction (Lika et al. 2011a, b). Here, we chose to evaluate variations around the lower value of *κ* to represent more realistic scenarios where limiting food can alter the reproductive output (Lika and Kooijman 2003). Thus, we assessed *κ* equal to 0.43, 0.51, and 0.58.

We consider the rest of the model parameters as constants (Table 2) because previous interspecific comparisons have shown that maintenance costs and structural costs remain largely similar between species (van der Veer et al. 2001, Freitas et al. 2010). To maintain biological consistency among all the parameters, we used the estimated values for *Daphnia magna*. The parameters of *D. magna* have been estimated from multiple experiments, reaching a data completeness of 6, which is the highest in the AmP collection. As such, these values are more likely to represent the individual physiology accurately.

**Table 2.**
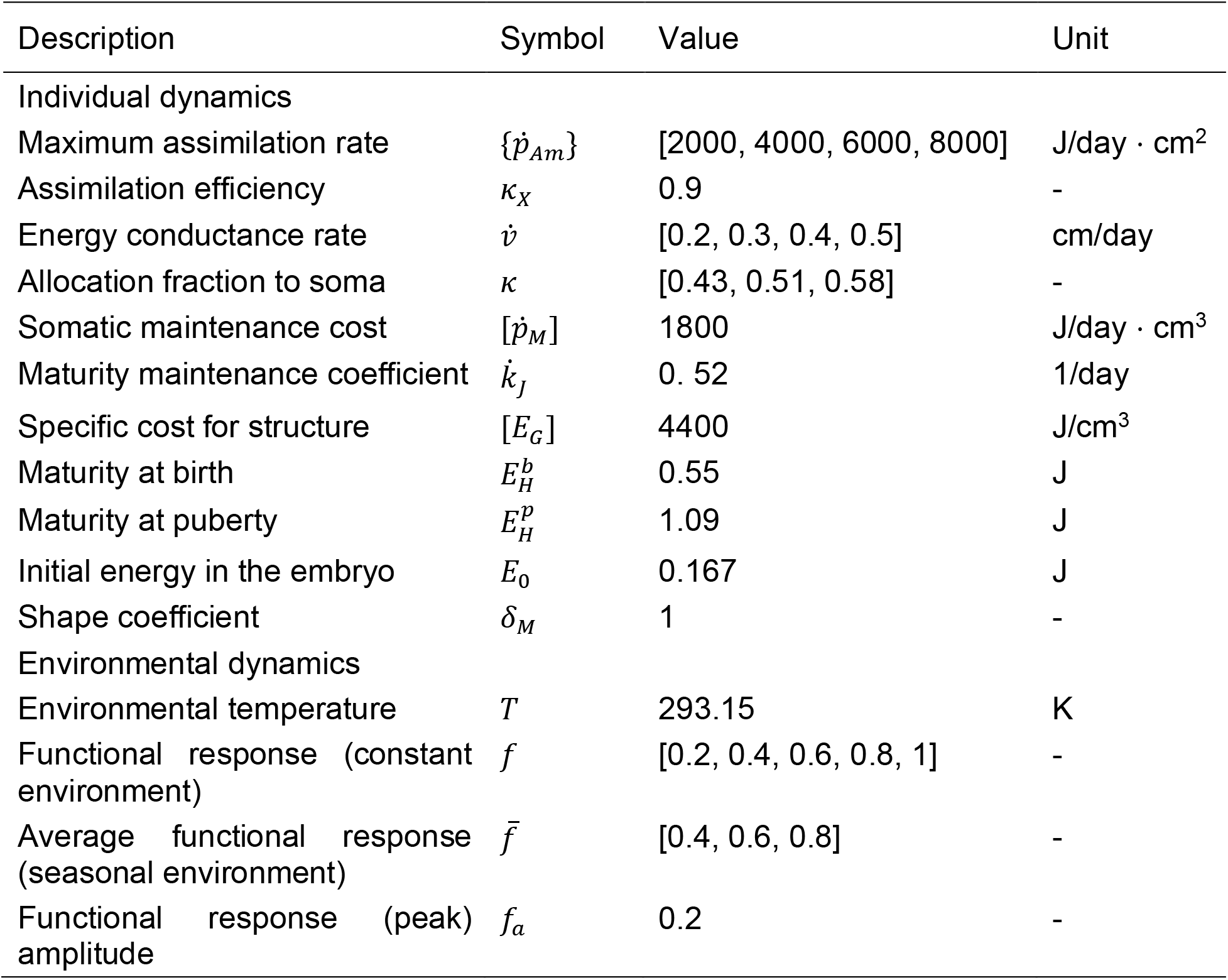
Parameter values for the simulations of the standard DEB model (equation 1) and the resource (equation 3). Notation: square braces ([]) indicate quantities related to structural volume, curly braces ({ }) denote quantities related to structural surface-area, dots (.) indicate rates.

| Description | Symbol | Value | Unit |
| --- | --- | --- | --- |
| Individual dynamics |  |  |  |
| Maximum assimilation rate | $\{\dot{p}_{Am}\}$ | [2000, 4000, 6000, 8000] | J/day · cm<sup>2</sup> |
| Assimilation efficiency | $\kappa_X$ | 0.9 | - |
| Energy conductance rate | $\dot{v}$ | [0.2, 0.3, 0.4, 0.5] | cm/day |
| Allocation fraction to soma | $\kappa$ | [0.43, 0.51, 0.58] | - |
| Somatic maintenance cost | $[\dot{p}_M]$ | 1800 | J/day · cm<sup>3</sup> |
| Maturity maintenance coefficient | $\dot{k}_J$ | 0.52 | 1/day |
| Specific cost for structure | $[E_G]$ | 4400 | J/cm<sup>3</sup> |
| Maturity at birth | $E_H^b$ | 0.55 | J |
| Maturity at puberty | $E_H^p$ | 1.09 | J |
| Initial energy in the embryo | $E_0$ | 0.167 | J |
| Shape coefficient | $\delta_M$ | 1 | - |
| Environmental dynamics |  |  |  |
| Environmental temperature | $T$ | 293.15 | K |
| Functional response (constant environment) | $f$ | [0.2, 0.4, 0.6, 0.8, 1] | - |
| Average functional response (seasonal environment) | $\bar{f}$ | [0.4, 0.6, 0.8] | - |
| Functional response (peak) amplitude | $f_a$ | 0.2 | - |

### Parameter space validation

To verify our parameter subset’s biological relevance, we examined the AmP collection for species within the parameter space. Accordingly, we used the routines in the AmPtool MATLAB package (AmPtool 2020) to find all the species parameterized with the standard DEB model within the limits of our parameter space for 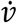 (in the interval [0.15, 0.55]), 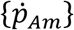 (in [1500, 9500]), and 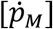 (in [1600, 2000]). We did not restrict the value of *κ* to our parameter subset because the collection is biased towards larger values. We quantified the parameter variation in these species through the coefficient of variation (*c_v_* = *σ/μ*).

### Relationship between variables and observable quantities

The state variables in the DEB model are not directly measurable; hence, we transform them into quantities that can be observed across individuals. Specifically, the reserve energy (*E*) plus the structural volume (*V*) constitute the energy fixed in the individual’s biomass. Nonetheless, to calculate the biomass as (dry) weight in grams, we need to account for the density and compositions of both variables, as given by:

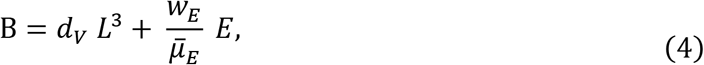

where *d_v_*, (g/cm^3^) is the density of the structural volume, *w_E_* (g/Cmol) is the molar weight of the energy reserve, and 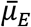 (J/Cmol) is the chemical potential of the energy reserve (equation 3.3 in Kooijman 2010). These constants are species-specific; for generality, however, we used the standard values: *d_v_* = 0.28, *w_E_* = 23.9, and 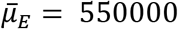 (Kooijman 2010).

We illustrated the model behaviour using the related Fan-tailed Gerygone, *Gerygone flavolateralis*, and the Grey Warbler, *G. igata*, as a case study to compare the differences between the constant and seasonal resource environments. For these two species, we transformed the above dry biomass (B) into wet biomass (B_*w*_) assuming the relation B_*w*_ = 5B, which is based on observations of water content in fledglings of the Grey Warbler (Gill 1982).

### Model analysis

When there are seasonal fluctuations, the model’s nonlinearities make it impossible to determine the dynamics analytically (Muller and Nisbet 2000). For this reason, we addressed our questions through numerical studies. We implemented the DEB model in the R language (version 3.6.2, R Core Team 2019), and performed the time integrations using the “lsoda” initial value problem solver from the package **deSolve** (Soetaert et al. 2010). We integrated the model for three years, and for the simulations considering resource variability, we assumed one seasonal cycle per year. However, we used an integration time of six years to compare the tropical and temperate species.

### Data analysis

To show the combined effect of the parameters and the resource, we summarized the model simulations at different food levels through scatterplots. We express the energy reserve and the structural volume variables together as the individual’s biomass for conciseness. For comparative purposes, in the constant resource environment, we plot the steady-state value of the biomass. In contrast, in the seasonal environment, we show the average values attained after transient dynamics have been discarded. For the cumulative reproduction energy variable, in both environmental scenarios, we show the average reproductive energy attained in the last two years of the individual’s lifespan. Similarly, for the maturity energy variable, we emphasize the time to reach the puberty threshold in both food environments.

To take into account the relative differences between individuals in our analysis, we computed the relative value of each state variable *x*:

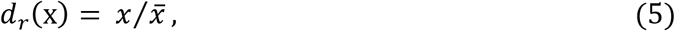

where 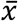 is the mean of all the simulations at the same resource level *f*, but considering a different parameter set for 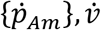, and *κ*.

### Model validation

At constant resource availability, the reserve equation in (1) leads to the constant:

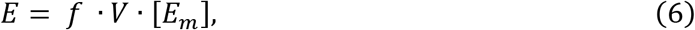

where 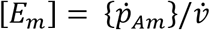, which is the maximum value of the reserve density (also called the reserve capacity). Thus, to verify our simulations’ behaviour, we compared our model predictions for energy reserves at constant resource with the expected values from equation 6.

## RESULTS

### Parameter space validation

We found 14 species in the AmP collection modelled with the DEB-std model having parameter values that lie within our parameter space for 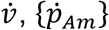, and 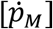 (Fig. 3, Table 3). These species’ parameters have values across all our parameter space for 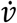; however, for 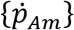 the values are below 8000 (Fig. 5). All of the species are in the Aves class (superorder Neognathae) likely due to the restriction of selecting entries parameterized with the DEB-std model (which constitute ~45% of the collection) and the high representation of this class in the collection (~21%).

**Figure 3.**
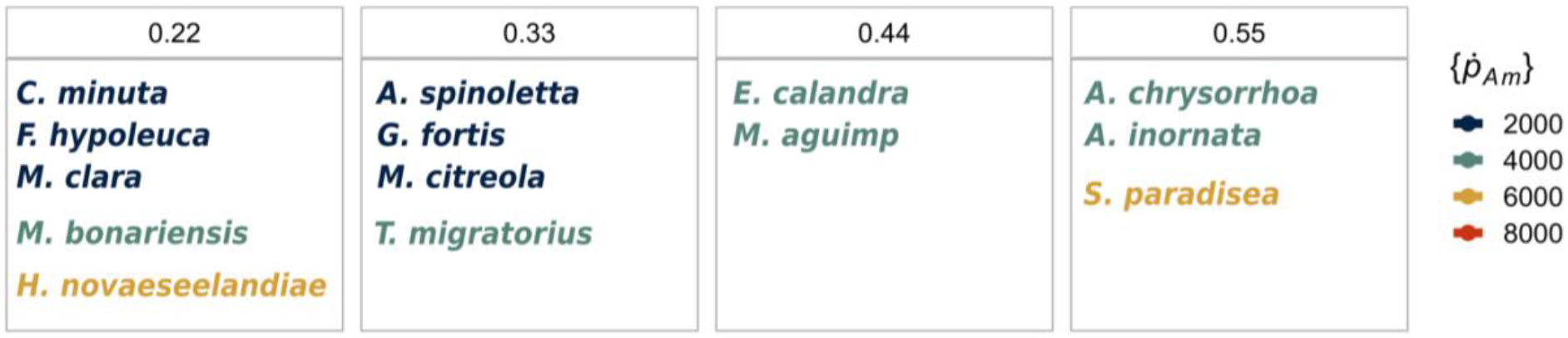
Position of the species within our parameter space for 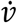 and 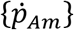. See Table 3 for parameter values and common names.

**Figure 4.**
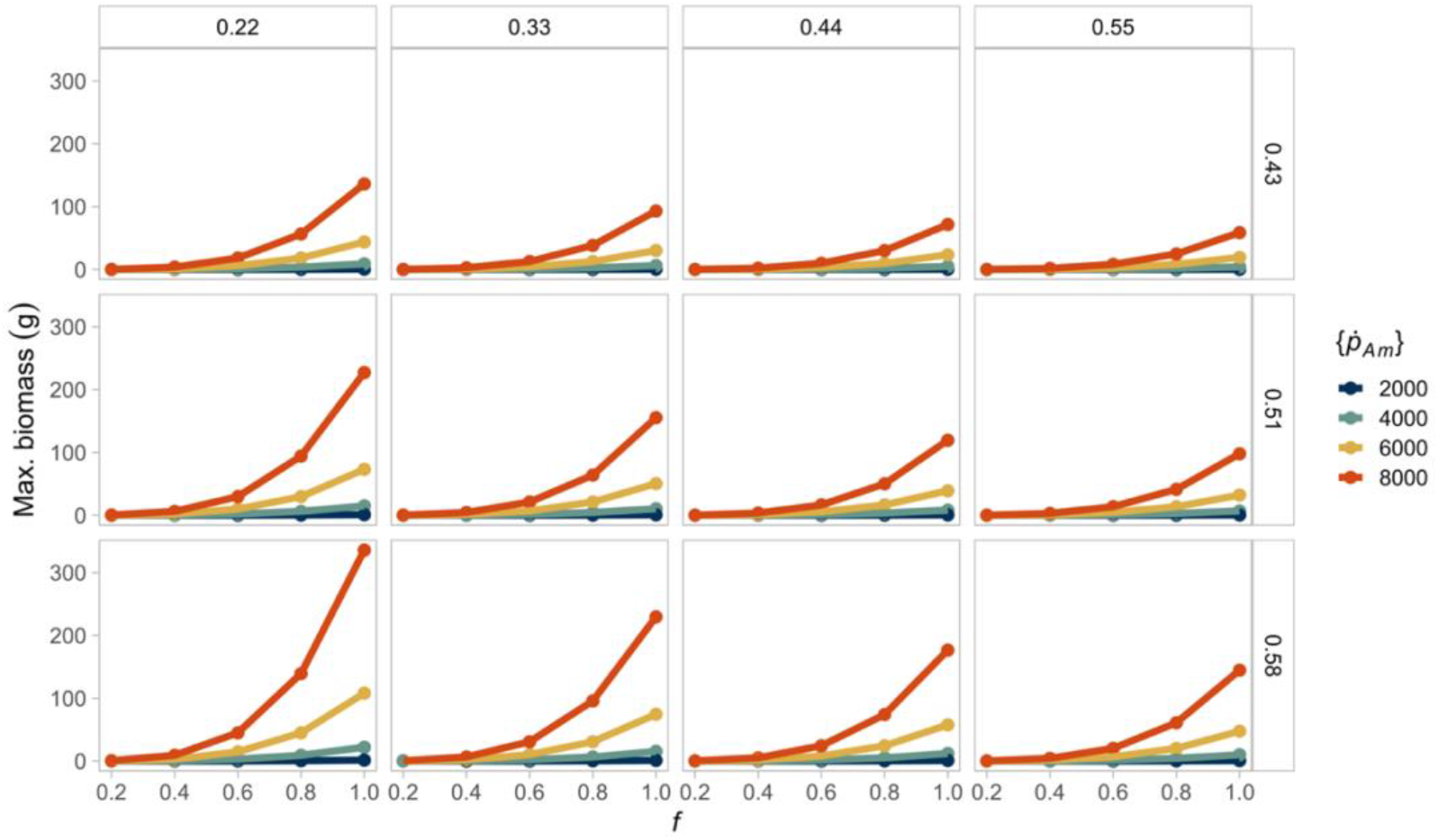
A decreasing, constant resource (*f*) reduces interspecific differences in maximum biomass. The largest biomass is attained when individuals combine high assimilation with low energy conductance. The columns show the different values of energy conductance evaluated 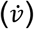. The rows represent the fraction of energy allocated to soma (*κ*). The colours of the lines indicate the value of the maximum specific assimilation rate 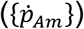. Lines of the same color in each box (equivalent to a parameter combination) represent the same species at different food levels.

**Figure 5.**
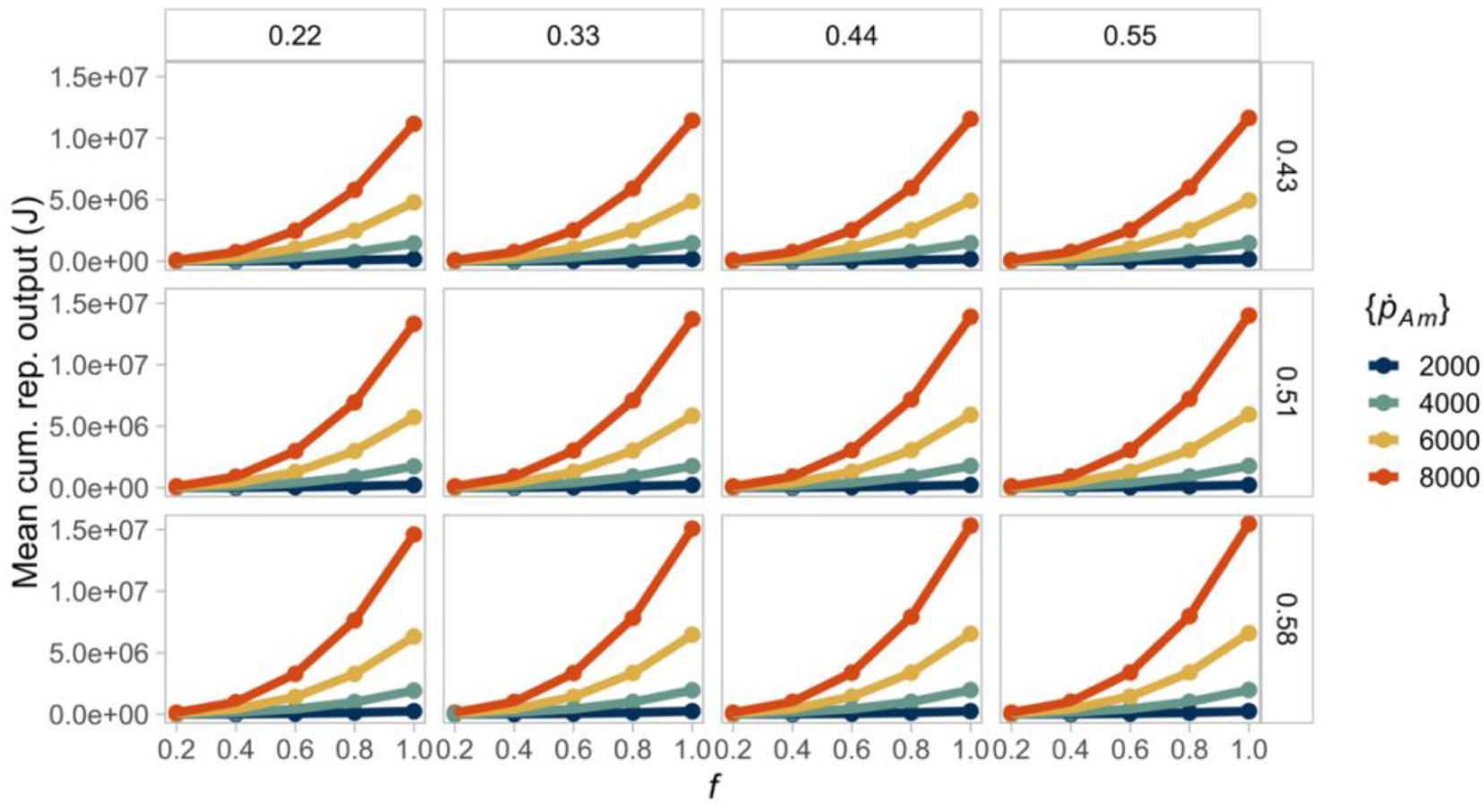
A declining, constant resource (*f*) reduces interspecific variability in mean cumulative reproductive output. Higher reproductive output is reached when the fraction of energy allocated to soma is high. The columns show the different values of energy conductance evaluated 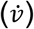. The rows represent the fraction of energy allocated to soma (*κ*). The colours of the lines indicate the value of the maximum specific assimilation rate 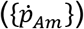. Lines of the same color in each box (equivalent to a parameter combination) represent the same species at different food levels.

The birds that fall within our parameter space are characterized by being small to medium size (from 15.5 g in the European pied flycatcher to 100 g in the Artic tern, except for the New Zealand pigeon at 590 g), terrestrial, flighted and mostly carnivores (mainly insectivores, except for the New Zealand pigeon and the Medium ground finch, which are frugivores or granivores). Relative to the AmP collection (AmP 2021), the species’ parameters are biased to high values of energy conductance and somatic maintenance costs and intermediate to high assimilation values. The parameters for [E_*G*_], 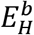, and 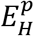, are all above the limits of our parameter space because they correspond to species of a larger size and longer lifespan than *D. magna*.

### Model validation

Our model’s predictions in the constant food environment match the analytical solutions for the reserve density (Fig. S1), which indicates that the simulations are consistent with the expected results.

### Effect of interspecific differences

To test for the consequences of interspecific variability, we conducted simulations in a constant resource environment. We expected that simulations where the resource is limiting produce similar size and reproductive output individuals because low food should reduce interspecific variability. We found that a decreasing resource does minimize the consequences of interspecific differences in biomass and reproduction (Figs. 4 and 5. See Figs. S1 and S2 for the differences in reserve energy and structural volume). We found the opposite effect for maturation time, where lower food levels lead to variable development rates (Fig. S4).

When the resource is non-limiting, a combination of high assimilation but low energy mobilization 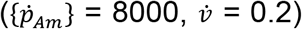 produces individuals with the highest biomass (Fig. 4). Here, a lower energy conductance magnifies the interspecific differences since the organisms mobilize less energy from the reserves. However, the fraction of energy allocated to soma does not affect the biomass because the increase in energy and volume is proportional.

Interspecific differences are greater in the reproductive output for organisms that combine a large assimilation rate with a high fraction of energy allocation to soma in an environment with high resource availability (Fig. 7). This counterintuitive result, where allocating more energy to soma produces higher reproduction (instead of lower reproduction), is caused by the moderate *κ* values in our parameter space: *κ* = 0.58 likely to be close to one of the two optimum points of maximum reproductive output as a function of *κ*. The energy conductance rate seems not to affect the reproductive output, which may be a consequence of the smaller range evaluated for conductance compared to that of energy assimilation.

**Figure 6.**
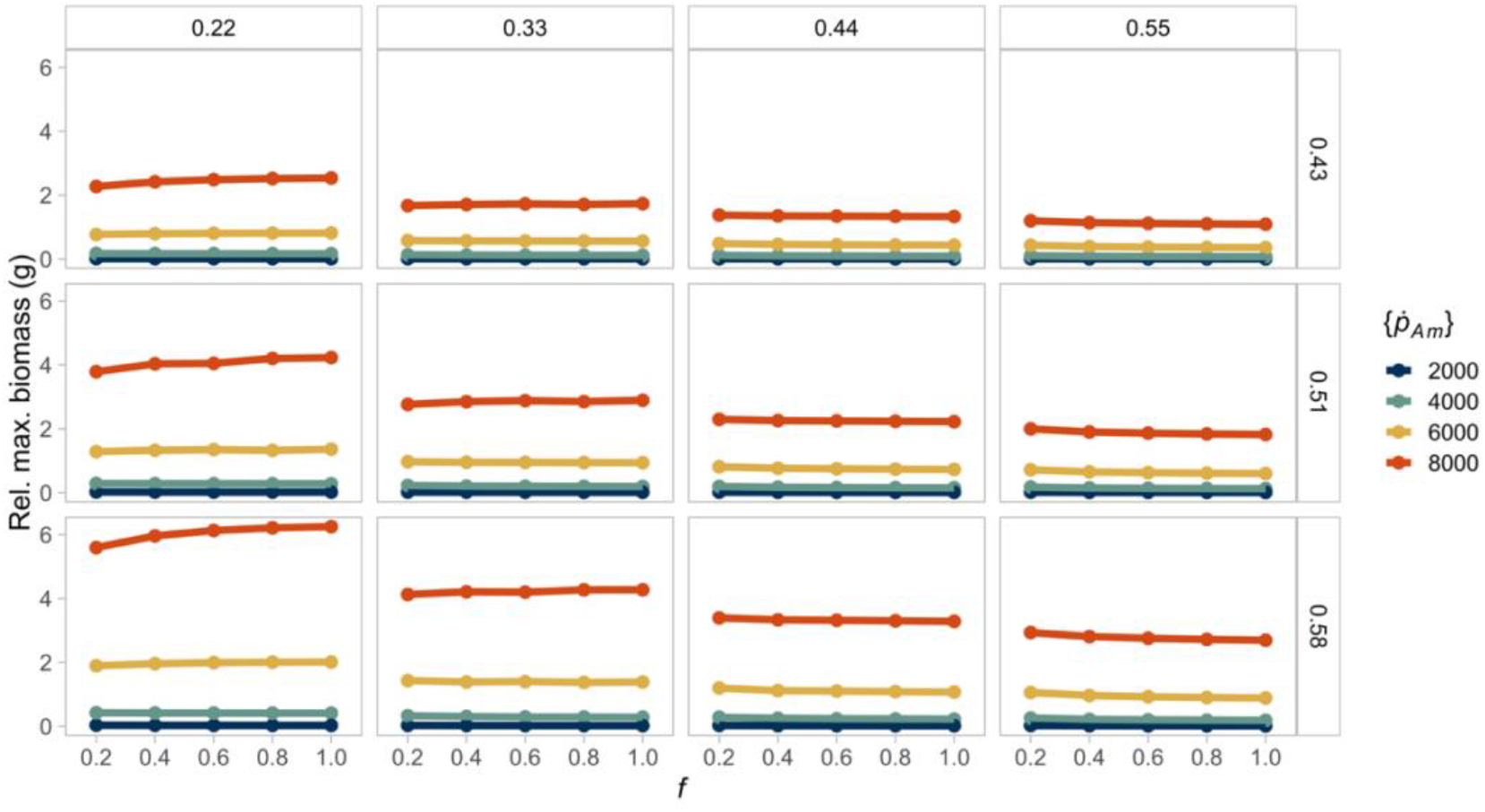
A constant resource (*f*) scales the interspecific differences in biomass. Hence, there are only small relative differences between different food levels for the same species. The columns show the different values of energy conductance evaluated 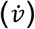. The rows represent the fraction of energy allocated to soma (*κ*). The colours of the lines indicate the value of the maximum specific assimilation rate 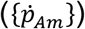. Lines of the same color in each box (equivalent to a parameter combination) represent the same species at different food levels.

**Figure 7.**
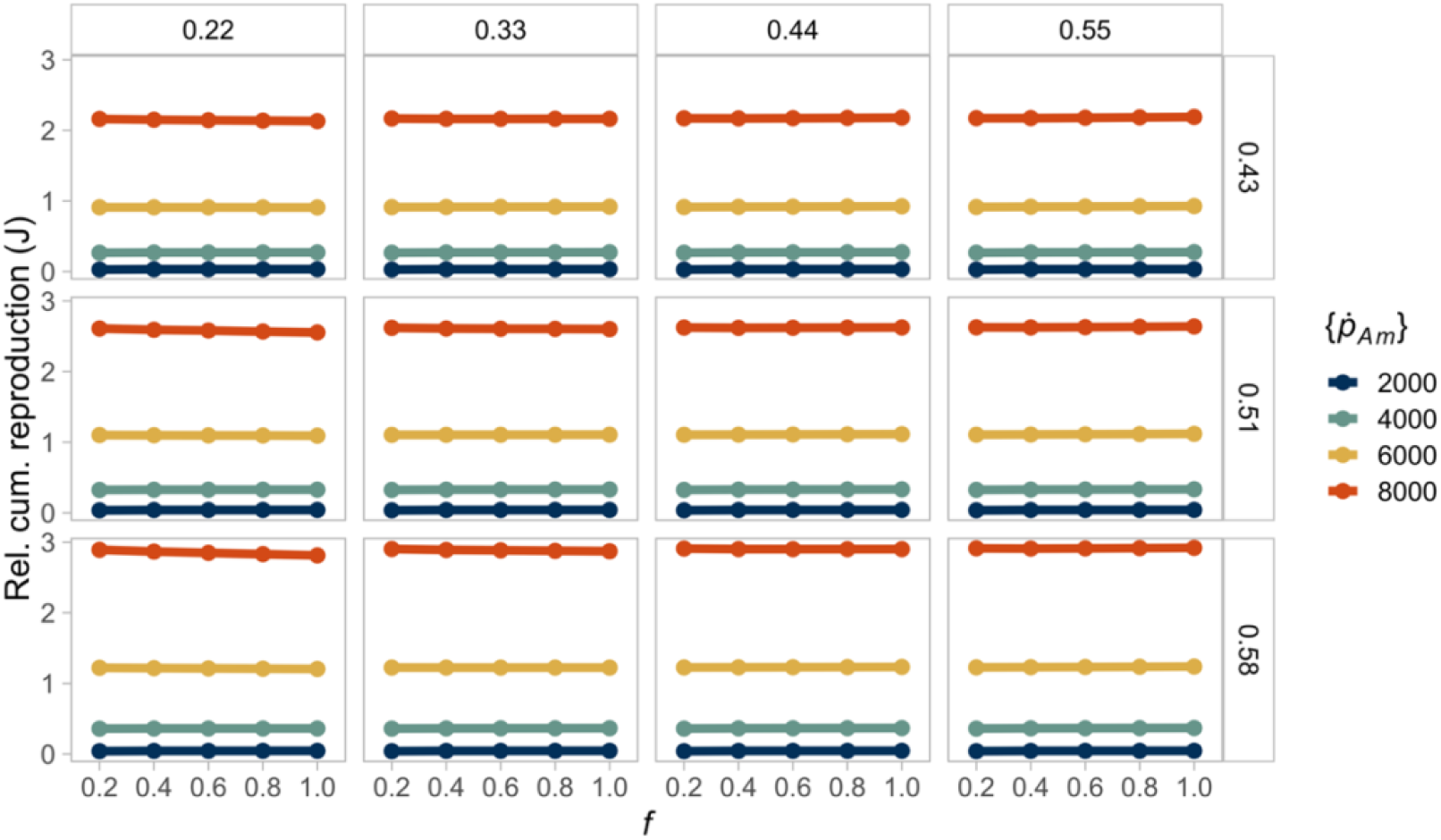
A constant resource (*f*) scales the interspecific differences in reproductive output. Hence, there are no relative differences between different food levels for the same species. The columns show the different values of energy conductance evaluated 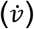. The rows represent the fraction of energy allocated to soma (*κ*). The colours of the lines indicate the value of the maximum specific assimilation rate 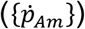. Lines of the same color in each box (equivalent to a parameter combination) represent the same species at different food levels.

The maturation time shows increased variability when the resource is scarce (Fig. S4). The differences in development time are small (between a minimum of 1 day to a maximum of 4 days); however, they are greater for individuals with reduced assimilation and conductance rates at an intermediate value of energy allocation to soma. For example, when food is scarce (*f* = 0.2), and the energy allocation to soma increases (*κ* = 0.58), a combination of low energy assimilation 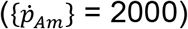 and mobilization 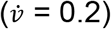 produces individuals with the slowest maturation (4 days).

Organisms grow larger at higher food, thus resulting in larger absolute differences. However, the relative differences in biomass (Fig. 6) are not strictly constant, with higher resource leading to relatively higher biomass when energy conductance is low 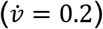 but to relatively lower biomass when mobilization is high 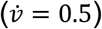. These differences in biomass reflect the effect of the energy conductance parameter on the energy reserves (Fig. S5), while the relative differences in structural volume remain constant (Fig. S6). Similarly, the relative differences in reproduction energy show that species’ differences are nearly constant across resource levels (Figs. 7 and S6) or, in the case of maturation time, the differences are small (Fig. S7). These findings indicate that the resource level mainly has a scaling effect on the individual dynamics.

For maturation energy, the relative differences are not identical for each resource level and appear to be larger for a combination of lower resource and energy conductance (Fig. S7). However, the range of these differences is small and may not reflect significant differences.

### Effect of resource variability

We evaluated the consequences of environmental variability by simulating a seasonally-varying resource. As in the environment with a constant resource, absolute interspecific variability in biomass and reproductive output is greater when average food is more abundant (Figs. S8 and S9). Further, biomass and cumulative reproductive output are independent of the initial resource density (Figs. S13 to S21), indicating that the individuals can compensate for variations in resource abundance during their lifespan. As expected, the average values for biomass and cumulative reproductive output are larger compared to the same individual in a constant environment with equal mean food availability. We address these results in the next section.

As in the constant environment, the maturation time shows increased variability when the resource is scarce (Fig. S12). Furthermore, interspecific differences are amplified when the initial resource density is low, compared to simulations where the initial resource is high. Individuals born in an environment that slowly becomes hospitable grow slower and remain small during the first season, particularly if they have a low assimilation rate. In contrast, individuals that start at the onset of a good period grow quickly to a large size, especially when the allocation fraction to soma is large. Yet, these differences in maturation may not be significant, given that the development times are equally short (Figs. S22 to S24).

### Comparing temperate and tropical species: an example

To better understand the resource’s effect, we compared the temporal dynamics of one species (given by the parameter set 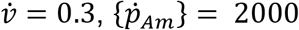, *κ* = 0.3, and 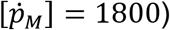 in both a constant and a seasonal resource environment with equal mean resource availability. The dynamics show that, despite having the same parameter values, the individual in the seasonal environment reaches a greater average biomass (Fig. 8A) and cumulative reproductive output (Fig. 8B) than an individual of the same species in a constant environment with an equal average resource. The resource does not appear to affect the maturation dynamics, given that the maturation threshold parameters are low and can be reached shortly after birth in both environments (Fig. 8C).

**Figure 8.**
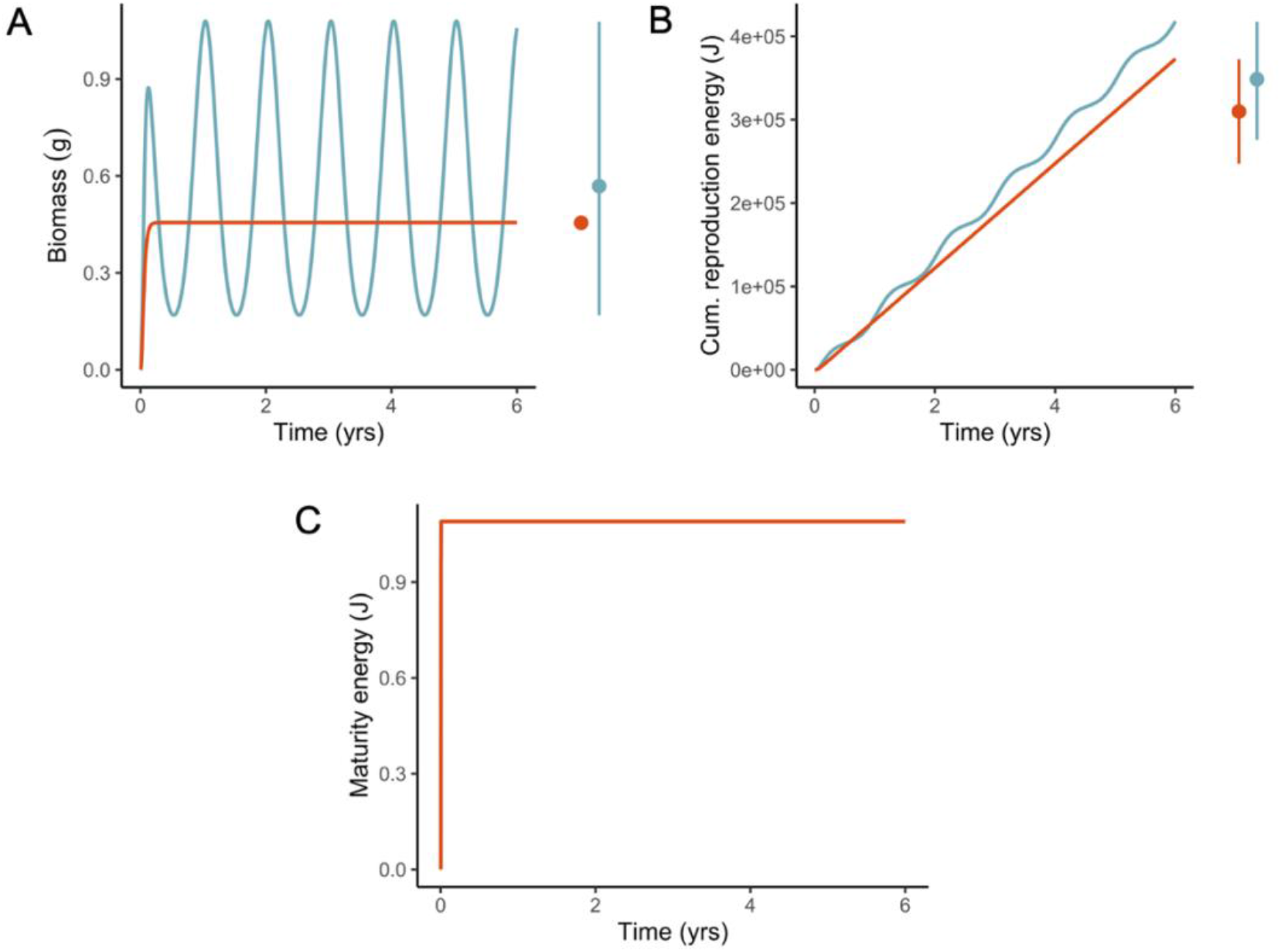
Individuals of the same species reach larger average biomass (A) and reproduction (B) in a seasonal resource environment (blue lines) relative to the same individual in an environment with an equal mean resource availability (red lines). In the right panel of (A), for the species in the constant environment, the red point shows the steady-state value of biomass reached at the end of the lifespan. In contrast, for the seasonal environment, the blue point represents the average biomass calculated over the last four years (i.e., years two to six), and lines show the minimum and maximum values. In the right panel of (B), for both species, points represent the average cumulative reproduction energy calculated over the last four years, and lines correspond to the minimum and maximum values. The biomass of the individual in the seasonal environment fluctuates according to the resource because the standard DEB model does not consider limits to individuals’ shrinking in size in periods of low food availability. The maturation dynamics (C) are not affected by the resource because the maturation energy threshold parameters are low, and individuals reach puberty shortly after birth. For the simulations, we assume 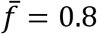, and the parameter set of the species is: 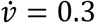, 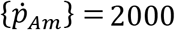, *κ* = 0.3, and 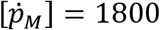 (see other parameter values in Table 3).

We further illustrate the model behaviour for the Fan-tailed Gerygone, *Gerygone flavolateralis*, and the Grey Warbler, *G. igata*, two related species with parameter values close to each other (Fig. 5, Tab. S2). Members of the *Gerygone* genus (Passeriformes, Acanthizidae) are small insectivorous species distributed in the Australasian region (Keast and Recher 2001), which do not migrate. The Grey Warbler inhabits temperate forests in the South Island of New Zealand, while the Fan-tailed Gerygone dwells in the tropical rainforests and savannahs of New Caledonia and Vanatu (Gill 1982, Attisano et al. 2019). The species differ slightly in their biomass: 6.45 g for the Grey Warbler (Gill 1982) and 6.1 g for the Fan-tailed Gerygone (Attisano et al. 2019). The differences in the reproductive output are more pronounced: the Grey Warbler has two broods per year with an average clutch size of four eggs (Gill 1982); in contrast, the Fan-tailed Gerygone has only one brood per year with a mean of two eggs (Attisano et al. 2019).

As expected, our simulations show that, with similar average food density, the Grey Warbler can reach a similar size and have a greater reproductive output relative to the Fan-tailed Gerygone (Fig. 9). These differences in reproduction are more pronounced than in the previous example (Fig. 8) because the species differ in their parameter values, mainly on the energy allocated to soma and the cumulative energy at birth and puberty (Tab. S2). Furthermore, the dynamics of the Grey Warbler show pronounced seasonal fluctuations because the standard DEB model does not consider limits to individuals’ shrinking in size in periods of low food availability. Consequently, the biomass and reproductive energy of the Grey Warbler fluctuate according to resource availability. Our predictions underestimate both species’ average biomass likely because we do not consider that the reproductive energy contributes to the overall weight of the individuals. This assumption allowed us to measure a continuous reproductive output across our simulations, but it may result in biomass underestimation in species that store reproductive energy between discrete reproductive seasons, such as the Grey Warbler or the Fan-tailed Gerygone. Moreover, we assume that the effect of temperature on metabolic rates is negligible. Despite these simplifying assumptions, our forecasts agree with the general pattern between the species and demonstrate the model’s behaviour as well as the importance of the resource in contributing to the overall dynamics.

**Figure 9.**
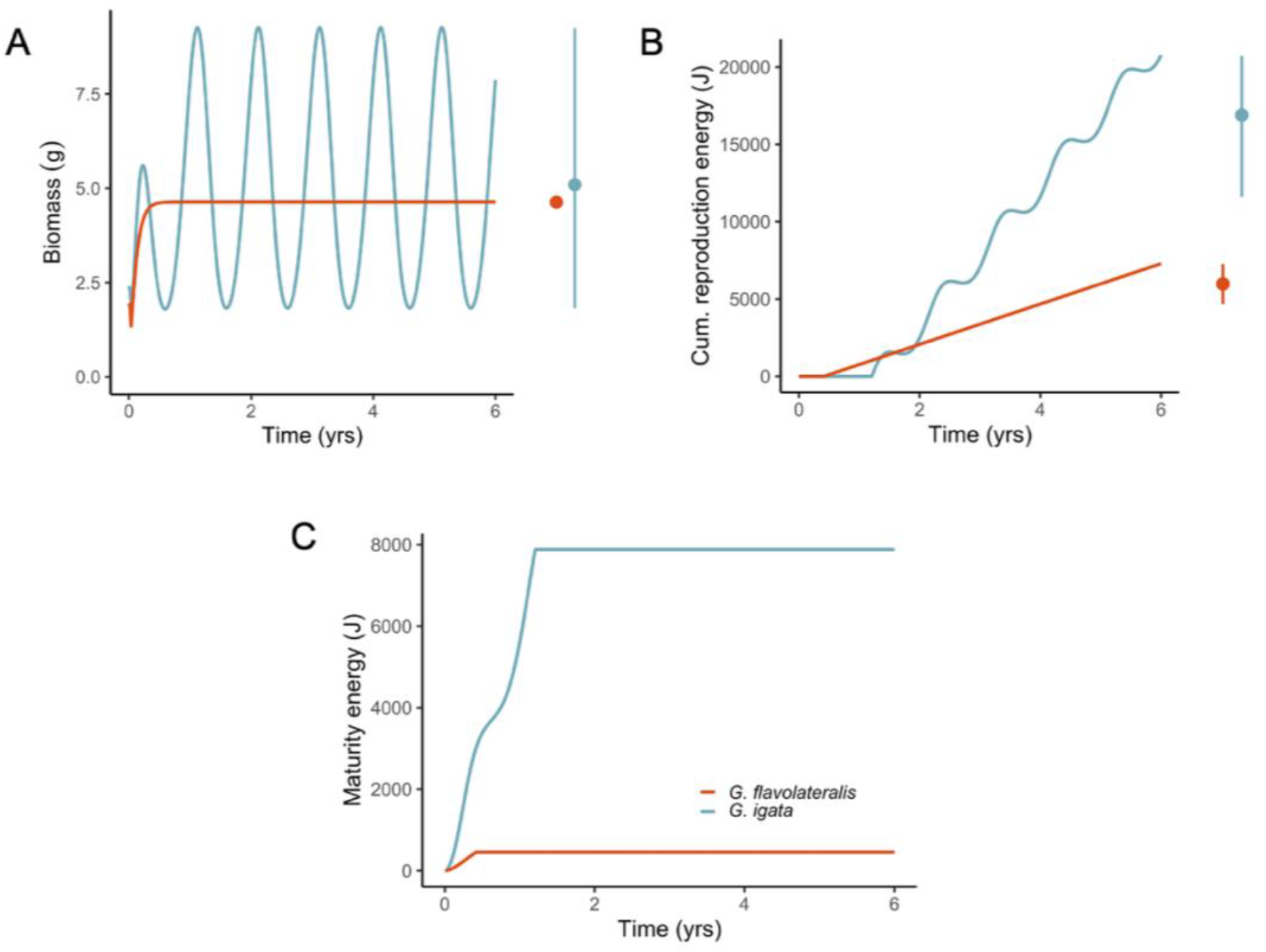
Simulations for the Grey Warbler (*G. igata*) in a seasonal environment with an equal mean resource availability as the Fan-tailed Gerygone (*G. flavolateralis*) in a constant environment show that the Grey Warbler dynamics follow the resource oscillations and reach a greater average wet biomass (A) and reproductive output (B) compared to the Fan-tailed Gerygone. In the right panel of (A), for the species in the constant environment, points show the steady-state value of wet biomass reached at the end of the lifespan. In contrast, for the seasonal environment, points represent the average wet biomass calculated over the last four years (i.e., years two to six), and lines show the minimum and maximum values. In the right panel of (B), for both species, points represent the average cumulative reproduction energy calculated over the last four years, and lines correspond to the minimum and maximum values. The Grey Warbler biomass fluctuates according to the resource because the standard DEB model does not consider limits to individuals’ shrinking in size in periods of low food availability. Differences in the maturation energy between the species (C) are due to different values of the puberty threshold (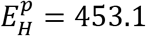 for the Fan-tailed Gerygone and 7880 for the Grey Warbler). The simulations assume 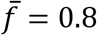 (see parameter values in Table S2).

## DISCUSSION

We used the DEB model to describe the individual rates of energy acquisition and partitioning, which lead to different life-history traits. Our approach is novel because we used simulations of the DEB model to perform a comparative analysis of 48 different strategies of energy allocation (equivalent to different species) across different environments to evaluate their consequences in body mass and reproductive output. As expected, we found that absolute interspecific differences in biomass and reproduction are more evident when the resource is non-limiting (Figs. 4 and 5). The same pattern holds in a seasonal resource environment; moreover, seasonality produces individuals with a greater average biomass and reproductive output relative to the constant environment (Figs. S8 and S9). Our results provide a plausible mechanistic explanation to known interspecific patterns of body size and reproductive output variation and reaffirm the importance of considering the effects of resource availability changes to predict broad biodiversity dynamics.

### Non-limiting resource amplifies interspecific variability in biomass and reproduction

Food availability per individual has been proposed as the primary cause of differences in adult body size and biomass, both among and within species, because it underlines organisms’ nutritional requirements, which ultimately drive ontogenetic growth (Huston and Wolverton 2011). Here, we showed that an abundant resource increases absolute interspecific variability in biomass, regardless of the environment (Fig. 4). However, species’ differences in biomass are not exactly constant, especially at low values of energy mobilization (Fig. 6). Although these differences are small, they indicate that the allometric relationships are not strictly proportional. Such disparities in biomass may suggest a larger sensitivity of the model output to parameter combinations with low mobilization and high assimilation at reduced food availability. Nevertheless, our simulations revealed that, in the DEB model, the resource has a scaling effect on the individual’s biomass because a larger food availability directly increases the feeding, assimilation, mobilization, and somatic maintenance fluxes, which result in a greater growth rate.

Our findings are in line with the central role of food availability on biomass (reflected in Bergmann’s rule, sizes in deserts, insular dwarfism, Dehnel’s phenomenon, and Cope’s rule, or more generally the resource rule), by which species become larger or smaller according to the size, abundance, and availability of food (McNab 2010). Although we did not include the effect of environmental temperature, several case studies using the DEB model have previously illustrated the relevance of food dynamics over temperature in determining growth rate and maximum possible size across different taxa (e.g., Cardoso et al. 2006, Freitas et al. 2009, Marn et al. 2019). For example, in two parapatric and genetically distinct populations of loggerhead turtles, a reduction in ultimate size has been shown to be a consequence of constant low food availability (Marn et al. 2019). Thus, within the DEB theory, a reduction in adult size or biomass can be interpreted as a direct consequence of the change in resource and its effect on assimilation (Kearney 2021), giving further theoretical support to the resource rule.

Animal body size is only one of the traits affected by resource availability; many other characteristics of individuals depend on the extent to which they can be afforded, including maintenance, reproductive output, and activity level (McNab 2010). Moreover, size alone can be considered a determinant of an organism’s ecological and physiological properties, such as reproduction (Klingenberg and Spence 1997, Litchman et al. 2013). Our results indicated that an abundant resource intensifies not only absolute interspecific variability in biomass in the DEB model, but also in reproductive output (Fig. 5). Our simulations showed that the resource has a similar effect on the individual’s reproductive output, where a greater food availability results in increased cumulative reproduction energy. The limiting effect of food quantity or quality in animal reproduction is well known in the literature (Kozlowski et al. 2020). Nevertheless, this conclusion is not always evident when making interspecific comparisons through the DEB model because of the covariation among parameters. For example, in the aforementioned loggerhead turtles, there is no significant disparity in the reproductive output of populations inhabiting areas with dissimilar food availabilities due to differences in the maturity maintenance and maturity energy thresholds parameters between individuals of the two populations (Marn et al. 2019). However, by fixing these parameters, we have isolated the scaling effect of resource availability on cumulative reproductive output.

In our formulation, we attribute interspecific variation to parameter values. Among the parameters that we evaluated, we found that both maximum assimilation 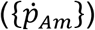 and allocation to soma (*κ*) have the most important role in promoting interspecific differences in individuals’ size and reproduction (Figs. 4 and 5). Such effect has previously been recognized among several marine invertebrates and vertebrates, where species differ mostly in their assimilation rate and the energy partitioning to growth and reproduction (van der Veer et al. 2001, Freitas et al. 2010, Marn et al. 2019). Both are parameters that can be considered highly adaptative, and their values are expected to reflect the conditions of the environment where the species evolved (Muller and Nisbet 2000).

### Environmental variability increases average biomass and reproduction

Food availability is influenced by biotic and abiotic factors, which can in turn covary, causing resource oscillations in time and space that determine the geographical and temporal changes in body size (Yom-Tov and Geffen 2006, 2011). By fixing the environmental temperature and evaluating different fluctuating resource scenarios, we showed that the surplus in food availability during one season of the year produces organisms that can reach a greater average biomass relative to the same individual in an environment with an equal average but constant resource (Figs. 8A and 9A). Our results are in agreement with the expectation according to the eNPP rule, in which animals subjected to fluctuations in food availability can do more than compensate for the periods of food deficit during their lifespan, as they gain more body mass during the periods of food surplus (Geist 1987, Huston and Wolverton 2011). In the DEB framework, it has been reported that the maximum size rises with the period and amplitude of the resource cycles, rather than to the mean (Muller and Nisbet 2000). Thus, our findings are in line with the DEB literature and give further support that indicates that the DEB model offers a mechanistic explanation at the interspecific level for the eNPP rule.

As proposed in the eNPP rule, food fluctuations in quantity and quality have further implications in traits associated with evolutionary fitness, most notably reproduction (Huston and Wolverton 2011). In our simulations in a seasonal environment, we found that the peaks in food availability also produce animals with a larger cumulative reproductive output compared to the same individual in an environment that has an equal average constant resource (Figs. 8B and 9B). Our results are consistent with previous studies that mention an increase in brood size in birds with latitude (Lack’s rule) or, more broadly, eNPP (Boyer et al. 2010). Nonetheless, because of the non-linearities that arise in the DEB model when there are food fluctuations, it has been shown that the total reproductive energy is highly dependent on the organism’s energy partitioning as given by the parameter values (Muller and Nisbet 2000). For example, a decrease in the value of *κ* (the energy allocation to soma) implies an increased commitment of energy to reproduction, and would be expected to simultaneously decrease size, shorten development and maturation times and increase reproduction allocation (Kearney 2021). However, as our results illustrate, individuals can also have a higher reproductive output when they allocate a greater fraction of energy to soma. Such behaviour occurs because maximum reproductive output as a function of *κ* has two optimum values: at intermediate (*κ* ≈ 0.5) and also at high values (*κ* ≈ 0.9) (Lika et al. 2011b). Hence, for the parameter combinations that we assessed, *κ* = 0.58 is likely to correspond to that intermediate optimum value that yields a greater reproductive output. Given that we simulated a restricted range of *κ*, we would expect that higher values (0.58 < *κ* ≲ 0.8) produce individuals for which reproduction is reduced because an increase in the energy allocation to soma will also increase the maintenance requirements (Muller and Nisbet 2000). The pattern of greater reproductive output for increasing values of *κ* also occurs in the seasonal environments that we evaluated. It has been shown that organisms with low to intermediate values of *κ*, such as the ones we simulated, reproduce more as the amplitude in food fluctuations increase (Muller and Nisbet 2000). Thus, depending on the underlying life history of the individual, resource seasonality in the DEB model may increase the reproductive output.

### Resource availability may influence development rates

According to the eNPP rule, the highest ontogenetic growth rates will occur where food availability is highest (Huston and Wolverton 2011). Consequently, when the resource is limiting, a delay in the maturation and reproduction rates is expected. For example, in two sympatric and sibling insectivorous bat species (genus *Myotis*) with marked differences in resource supply, there is a delay in the reproduction onset for the species with a lower food availability; however, this difference disappears in years where there is a pulsed input of a secondary resource (Arlettaz et al. 2017). In general, our results suggest that there is a delay in maturation time at lower, constant resource levels (Fig. S4). In a seasonal environment, our findings indicate that individuals that start their life at different points of the seasonal cycle do not have the same maturation times (Figs. S22 to S24).; More specifically, individuals born during the low food season show a slower development and more variable times to reach puberty, compared to the same organisms born during the high food season (Fig. S12). Similar results have been described for mussels parameterized with the DEB model (Muller and Nisbet 2000), highlighting the relevance of birth timing in developmental times when resource oscillates. Nevertheless, in both environmental scenarios, our results may not represent a significant difference as the range of the developmental time variation is small because the parameters that define the maturation thresholds are close to each other and remain constant. Thus, even though we find indications of a possible effect of food level on maturation time, further analyses are needed to reveal the potential variation on development according to the resource in the DEB framework.

### Life-history traits and parameter space

By matching our parameter subset to real species, we can explain various patterns within the parameter space. We found that the species in our parameter subset combine an intermediate assimilation rate 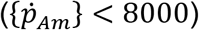 with energy conductance ranging from intermediate to high values (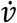 in the interval [0.15, 0.55]), which results in small body weight. In birds, such combination of values for energy conductance and assimilation have been associated with flying adaptations since it usually implies a low reserve density (i.e., a smaller contribution of the energy reserve to biomass; Lika et al. 2014, 2019, Teixeira 2016, Augustine et al. 2019). Additionally, most of the species within our parameter space have a carnivore diet, which has been linked to larger assimilation and somatic maintenance rates given that it requires greater enzymatic attack and thick stomach muscles (Ricklefs 1996, Battley and Piersma 2005, Teixeira 2016). Only one species, the New Zealand pigeon, constitutes an exception to the previous pattern, as it is larger and primarily frugivorous. This species has an intermediate assimilation rate combined with a lower mobilization rate, which results in a greater biomass due to a larger energy reserve. In general, our subset of species reflects previous findings that indicate that the mobilization rate is one of the most variable DEB parameters for birds (Teixeira 2016), and seem to suggest a broad link between assimilation rate and diet.

In birds, the average fraction of energy allocated to soma is very high (with a mean of 0.988) and does not exhibit significant variation among species (*c_v_* = 0.03 Teixeira 2016). This evidence suggests that the energy allocated to soma is phylogenetically conserved and that selective pressures have mostly driven birds towards larger investments in growth and maintenance, as well as delayed maturation and relatively low production efficiency (Teixeira 2016, van der Meer et al. 2020). In our species subset, *κ* ranges from 0.822 to 0.999, and the combination with intermediate assimilation and high somatic maintenance in some species could indicate that they have evolved greater maintenance costs that allow a fast growth to a small size, in accordance with the waste-to-hurry hypothesis (Kooijman 2013). Nonetheless, in birds, this combination of parameter values seems to be related to flight adaptations (Augustine et al. 2019) rather than to strategies that promote faster growth.

Traits associated with differing life histories are usually classified along a “fast to slow” or “pace-of-life” continuum. For example, compared to temperate birds, tropical birds are typically considered as having a “slow” life history, involving small clutch sizes or low annual reproductive output (Moreau 1944, Kulesza 1990), as well as slow growth and maturation of nestlings (Ricklefs 1968, 1976, 2000, Cox and Martin 2009, Martin et al. 2011, Jimenez et al. 2014). The results of our simulations comparing species in two resource environments agree with the literature, with the species in the seasonal habitat showing a faster growth to a larger body mass and greater reproductive output than the species in the constant habitat (Figs. 8 and 9). These differences between tropical and temperate birds have been attributed to trade-offs between investment in either reproduction or maintenance, as mediated by the biotic and abiotic environment (Roff 1992), which correspond to the species’ different parameter values. Nevertheless, our simulations also highlight the effect of the resource in modulating the species traits, which, together with predation risk, has been reported as the main environmental factor affecting growth rate and body size in birds (Bryant and Hails 1983, Pacheco et al. 2010, Martin et al. 2011, Jimenez et al. 2014).

### Model limitations

The DEB theory supposes that the parameters do not change over the individuals’ lifespan. Consequently, organisms show a passive phenotypic flexibility in response to resource availability, but there is no plasticity in the physiological mechanisms responsible for development and growth as a response to environmental cues (i.e., active adaptive plasticity). To modify this assumption would require discerning all possible changes in the conditions experienced by an organism during ontogeny (Pecquerie and Lika 2017). However, the current lack of understanding of how parameters may vary during an organism’s lifespan limits incorporating such responses into the DEB model (Freitas et al. 2009). Thus, to include an adaptive response in the individual energy allocation would require detailed knowledge of the species’ long-term seasonal patterns of growth and reproduction.

Quantitative and qualitative changes in food conditions would also be expected to affect individual survivorship and offspring production rate (Huston and Wolverton 2011). Simulations of the DEB model in a constant environment have suggested an increased lifespan as the resource becomes more abundant (Lika and Kooijman 2003). When food fluctuates, simulations have shown that individual lifespan is reduced as the oscillations amplitude increase (Muller and Nisbet 2000). However, we did not consider how resource availability affects individual lifespan or aging because such variations in lifespan are thought to be most significant for large values of *κ* (Muller and Nisbet 2000, Lika and Kooijman 2003), which we did not include in our simulations. Regarding the reproduction rate, we have not specified how reproductive energy is transformed into quantity and quality of offspring, given that the optimal strategy will depend on the particular environment experienced by each individual and its effect on growth, survival, and reproduction (Brown et al. 1993). Further elaborations of our study could incorporate such details, specifying how energy is allocated to survival and production of offspring over the individual’s lifetime.

The standard DEB model does not make any assumptions regarding starvation (Kooijman 2010). For this reason, the individuals in our model can decrease in size to cover maintenance costs when the resource is limiting, and energy reserves are exhausted. This assumption holds for many species, including platyhelminths, molluscs, and mammals (Genoud 1988, Downing and Downing 1993, Wikelski and Thom 2000, Saló 2006). Yet, other species may respond to food limitation in different ways. Nevertheless, we do not simulate periods of prolonged resource depletion or scarcity that could lead to starvation. Hence, the results of our simulation are still applicable to broad scenarios of seasonal food variation. Understanding other organisms’ strategies to survive seasonality without a reduction in structural mass remains an important issue for future research.

We illustrated the consequences of a constant and a seasonal resource by simulating the dynamics of two bird species parameterized for the DEB model. The fluctuations in biomass and reproductive output observed in the simulations with seasonality result from the lack of assumptions regarding starvation. Given that such a decrease in biomass is larger than the shrinking that birds are actually able to tolerate without dying, our simulations for the Grey Warbler serve mainly to exemplify a possible consequence of a seasonal environment in an organism’s physiology. Our results broadly agreed with the general pattern: temperate species can reach a larger reproductive output and slightly greater biomass than the tropical species. However, our inference is hindered because we did not use real estimates of food availability from the species’ habitats, and we simplified the abiotic environment only to resource availability, not considering temperature or biotic interactions, for example. Nevertheless, our purpose was to broadly show the different trends between a constant and a seasonal environment in closely related species.

We have shown how a mechanistic approach based on individual energetics can support empirical evidence on cross-specific trait variation. However, the geographical context of such ecological and evolutionary processes is an important component needed to gain a complete understanding of species’ trait gradients (Blackburn and Hawkins 2004, Olalla-Tárraga et al. 2010). Furthermore, any interspecific analysis must consider the phylogenetic non-independence of the data, i.e., closely related species are more similar than distant species (Olalla-Tárraga et al. 2010). For these reasons, more complex models that integrate a more accurate multidimensional environment with comparative phylogenetic methods are necessary to make stronger inferences of these eco-evolutionary patterns and processes.

## CONCLUSIONS

Using the DEB model, we separated the effects of genetically-determined physiological traits (represented by DEB model parameters) and resource availability. By fixing the resource, we showed that relative trait differences between species in biomass and reproduction are greater when food is non-limiting. We found similar results for simulations in a seasonal environment; moreover, resource fluctuations increase the individuals’ average biomass and reproductive output. Our results have potential implications for species of economic interest in which there is a desire to increase the yield in either biomass or reproductive output relative to the food consumption. Furthermore, our findings are a relevant step in forecasting organisms’ responses to environmental change. For example, global climate change has been linked to differences in the timing of resource availability and the arrival of migratory species to their feeding grounds. Finally, our simulations offer mechanistic support for patterns of body-size variation between related species arising from epigenetic effects (i.e., Bergman’s and Lack’s rules, or more generally, the resource and eNPP rules).

## ACKNOWLEDGMENTS

We thank the members of the Theoretical Biology Laboratory at MUN for their constructive comments on the manuscript. JM was supported by a School of Graduate Studies Baseline Fellowship from Memorial University and by a Mitacs Globalink Research Award. SCD and AH received funding from the Natural Science and Engineering Research Council of Canada (NSERC Discovery Grants 2015-06548 and 2014-05413).

