## Supplementary Material for "Resource and seasonality drive interspecific variability in a Dynamic Energy Budget model"

### Parameter space validation

There are 14 species in the AmP collection (on March 2020) modelled with the DEB-std model with parameter values that lie within our parameter space for $\dot{v}$, $\{\dot{p}_{Am}\}$, and $[\dot{p}_M]$ (Table S1).

**Table S1.** Species in the AmP collection with parameter values within our parameter space for $\dot{v}$, $\{\dot{p}_{Am}\}$, and $[\dot{p}_M]$. The coefficient of variation ($c_v$) summarizes the parameter variation across species. See Table 2 in the main file for parameter notation and units. The two species highlighted in bold (the Fan-tailed Gerygone and the Grey Warbler) are used to compare tropical versus temperate species.

| Species | Common name | $\kappa$ | $\{\dot{p}_{Am}\}$ | $\dot{v}$ | $[\dot{p}_M]$ | $[E_G]$ | $E_H^b$ | $E_H^p$ |
|---|---|---|---|---|---|---|---|---|
| *Acanthiza chrysorrhoa* | Yellow-rumped thornbill | 0.996 | 4724.4 | 0.50 | 1992.8 | 7328.6 | 54.9 | 10830 |
| *Acanthiza inornata* | Western thornbill | 0.995 | 4005.6 | 0.47 | 1881.1 | 7310.6 | 76.9 | 7243 |
| *Anthus spinoletta* | Water pipit | 0.997 | 3286.2 | 0.27 | 1670.3 | 7316.1 | 36.6 | 1974 |
| *Calidris minuta* | Little stint | 0.932 | 3898.4 | 0.19 | 1973.6 | 7338.4 | 1354.0 | 387000 |
| *Emberiza calandra* | Corn bunting | 0.915 | 4797.9 | 0.41 | 1949.0 | 7307.3 | 1758.0 | 999500 |
| *Ficedula hypoleuca* | European pied flycatcher | 0.822 | 3350.8 | 0.22 | 1642.7 | 7310.7 | 1389.0 | 823600 |
| *Geospiza fortis* | Medium ground finch | 0.968 | 3146.2 | 0.32 | 1644.7 | 7316.3 | 325.9 | 165000 |

| | | | | | | | | |
|---|---|---|---|---|---|---|---|---|
| *Hemiphaga novaeseelandiae* | New Zealand pigeon | 0.990 | 6900.6 | 0.17 | 1605.2 | 7316.0 | 570.1 | 616800 |
| *Molothrus bonariensis* | Shiny cowbird | 0.961 | 4122.9 | 0.21 | 1740.8 | 7332.9 | 766.0 | 277700 |
| *Motacilla aguimp* | African wagtail | 0.999 | 3804.9 | 0.45 | 1670.3 | 7290.4 | 13.4 | 1957 |
| *Motacilla citreola* | Citrine wagtail | 0.999 | 3204.1 | 0.31 | 1670.3 | 7312.9 | 6.4 | 860 |
| *Motacilla clara* | Mountain wagtail | 0.998 | 3109.4 | 0.22 | 1670.3 | 7330.5 | 15.1 | 1355 |
| *Sterna paradisaea* | Arctic tern | 0.991 | 5604.5 | 0.45 | 1681.2 | 7359.8 | 691.8 | 230000 |
| *Turdus migratorius* | American robin | 0.981 | 4940.2 | 0.32 | 1734.4 | 7306.1 | 440.3 | 312100 |
| Mean | | 0.967 | 4206.9 | 0.32 | 1751.9 | 7319.8 | 535.6 | 273994 |
| $c_v$ | | 0.05 | 0.26 | 0.36 | 0.08 | 0.00 | 1.10 | 1.20 |

**Model validation**

At constant food levels, the relative differences between our simulations for reserve density and the analytical solutions (equation 6) are close to zero (Figure S1). These small differences at each of the five constant resource levels we evaluated show the consistency of our simulations.

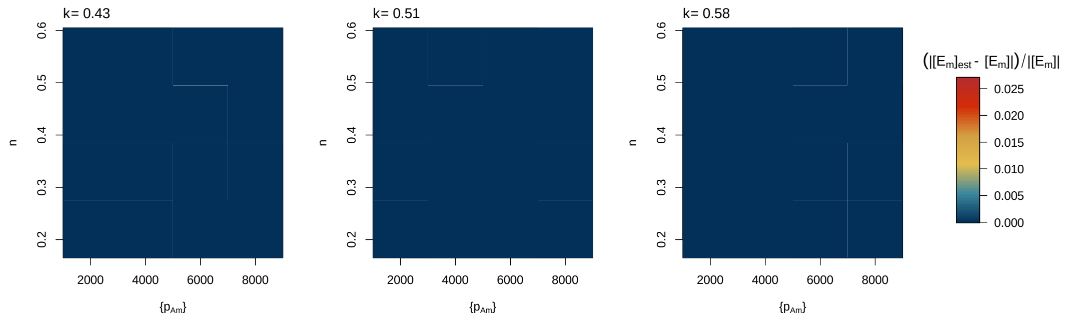
k= 0.43
k= 0.51
k= 0.58
n
0.2 0.3 0.4 0.5 0.6
2000 4000 6000 8000
{pAm}
(|[Em]est - [Em]|)/|[Em]|
0.025 0.020 0.015 0.010 0.005 0.000

27

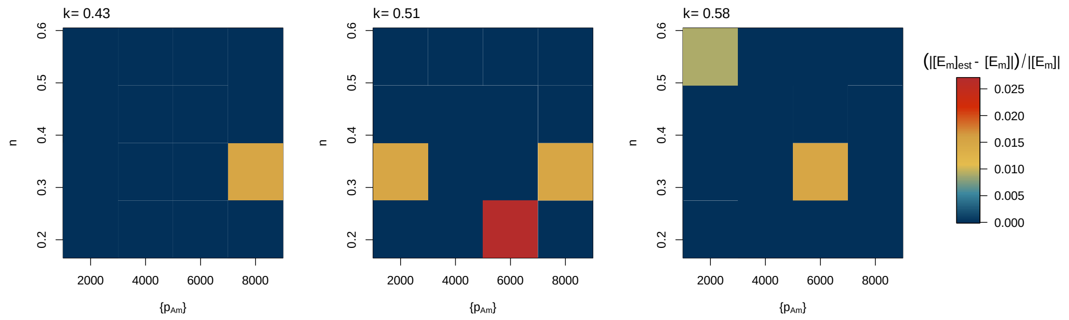
k= 0.43
k= 0.51
k= 0.58
n
0.2 0.3 0.4 0.5 0.6
2000 4000 6000 8000
{pAm}
(|[Em]est - [Em]|)/|[Em]|
0.025 0.020 0.015 0.010 0.005 0.000

28

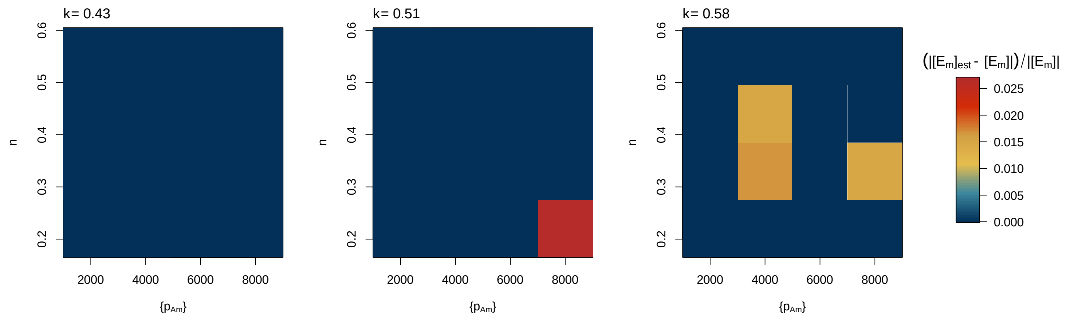
k= 0.43
k= 0.51
k= 0.58
n
0.2 0.3 0.4 0.5 0.6
2000 4000 6000 8000
{pAm}
(|[Em]est - [Em]|)/|[Em]|
0.025 0.020 0.015 0.010 0.005 0.000

29

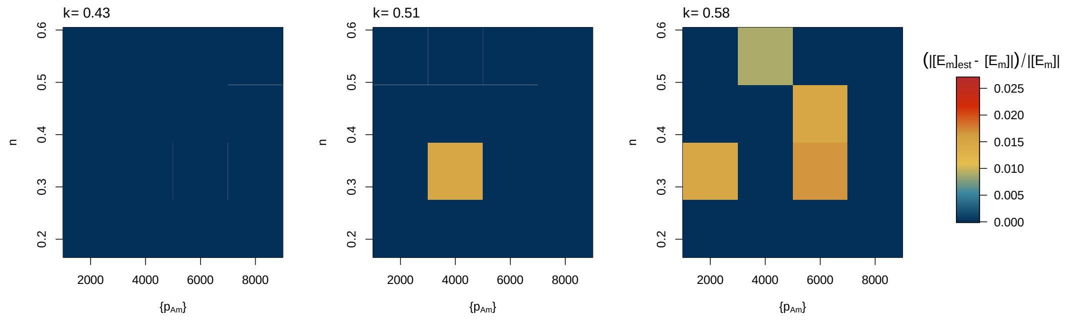
k= 0.43
k= 0.51
k= 0.58
n
0.2 0.3 0.4 0.5 0.6
2000 4000 6000 8000
{pAm}
(|[Em]est - [Em]|)/|[Em]|
0.025 0.020 0.015 0.010 0.005 0.000

30

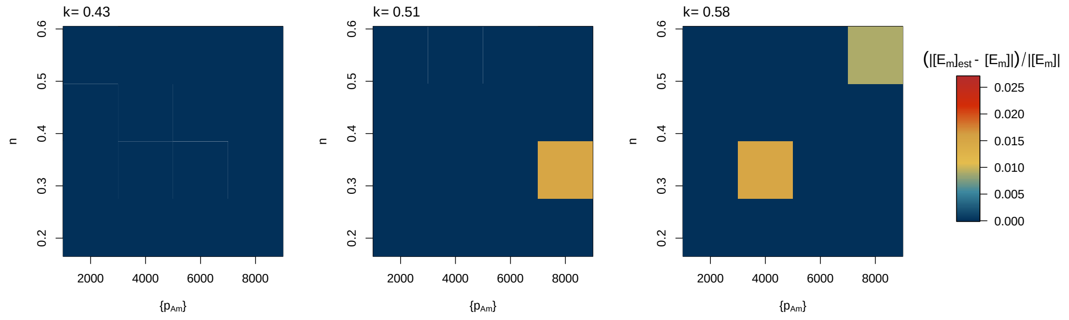
k= 0.43
k= 0.51
k= 0.58
n
0.2 0.3 0.4 0.5 0.6
2000 4000 6000 8000
{pAm}
(|[Em]est - [Em]|)/|[Em]|
0.025 0.020 0.015 0.010 0.005 0.000

31

**Figure S1**. The relative difference between the model simulations for reserve density ($[E_m]_{est}$) and the analytical solutions ($[E_m]$, equation 6) at a constant resource availability are close to zero. The rows correspond to the functional response at the five levels considered in descending order, i.e., from $f = 1$ to $f = 0.2$. The heatmap colour indicates the relative error between the numerical and the analytical solutions.

**Effect of interspecific differences**

When the resource is constant, a decreasing level minimizes the consequences of interspecific differences in reserve energy and structural volume (Figs. S2 and S3), which are directly reflected in the individual's biomass (Fig. 5). The effect is the opposite for the development time (Fig. S4), where the interspecific differences become greater as the resource decreases. However, the differences in time to reach puberty are small, ranging from 2 to 4 days; and are likely not significant.

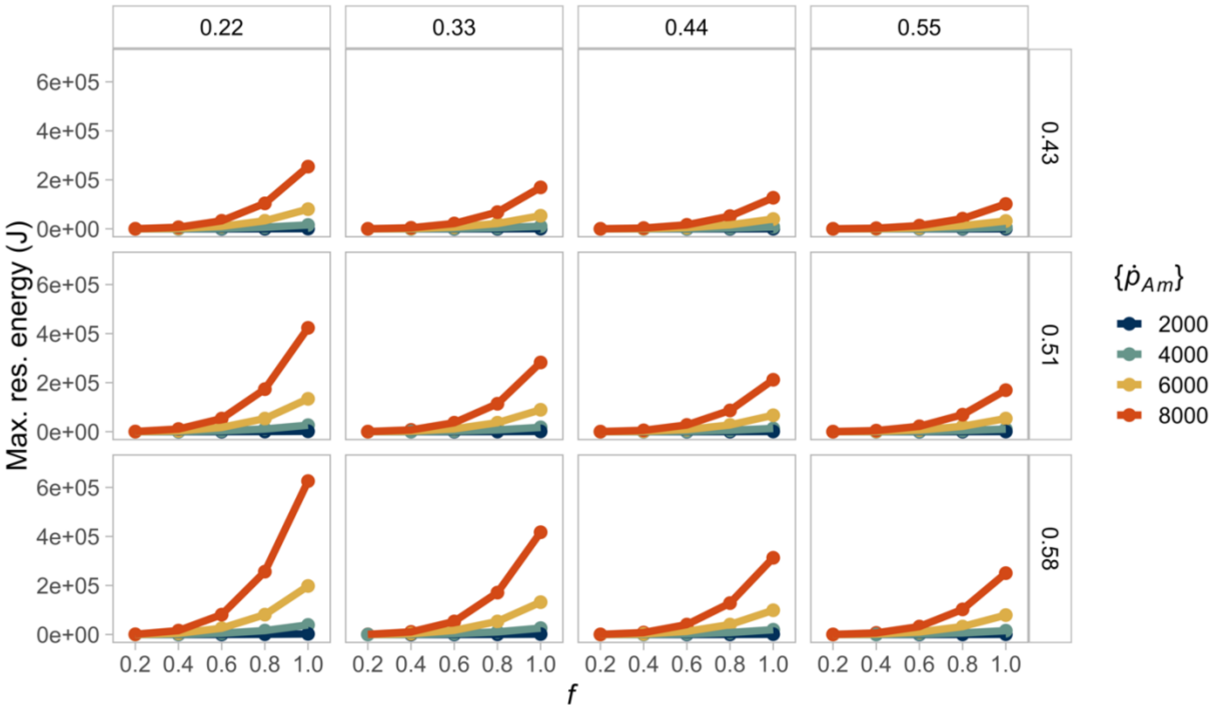


**Figure S2.** A constant, decreasing resource reduces interspecific differences in reserve energy. The largest reserve is attained when individuals combine high assimilation with low energy conductance. The columns indicate the value of energy conductance, while the rows represent the fraction of energy allocated to soma. Point and line colours indicate the maximum specific assimilation rate value. Points and lines of the same colour in each box (equivalent to a parameter combination) represent the same species at different food conditions.

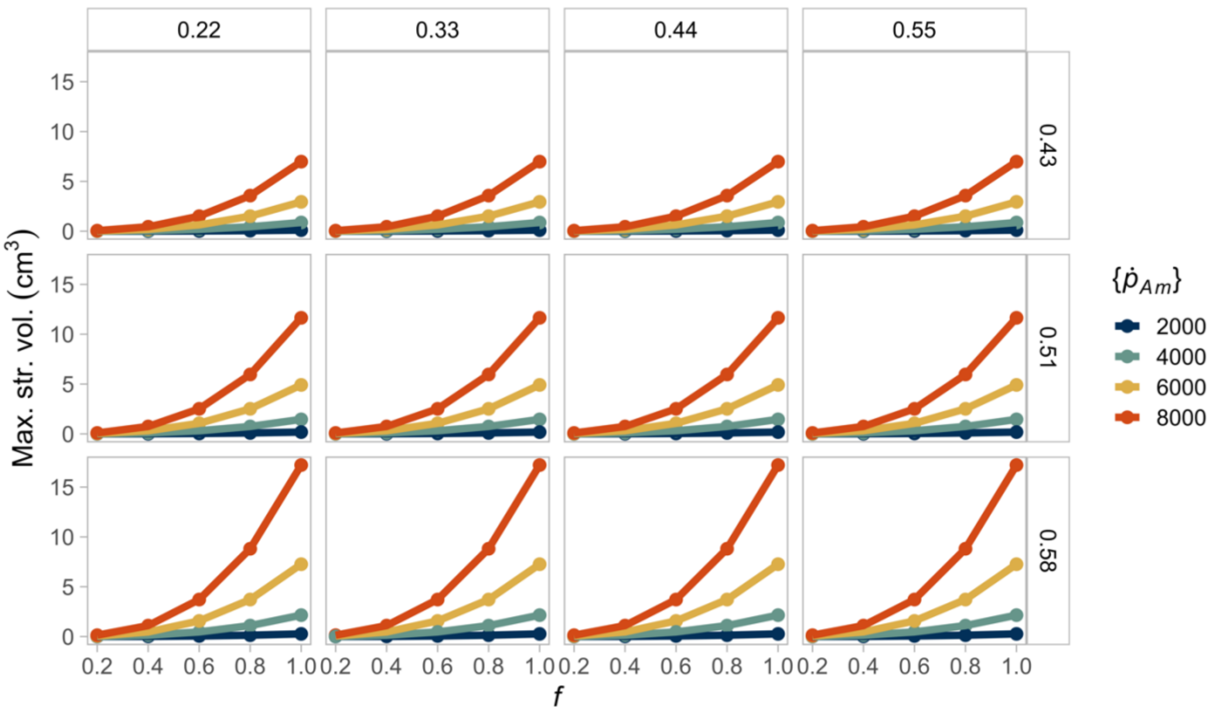


**Figure S3.** A constant, decreasing resource reduces interspecific differences in structural volume. The largest volume is attained when individuals combine high assimilation with low energy conductance. The columns indicate the value of energy conductance, while the rows represent the fraction of energy allocated to soma. Point and line colours indicate the maximum specific assimilation rate value. Points and lines of the same colour in each box (equivalent to a parameter combination) represent the same species at different food conditions.

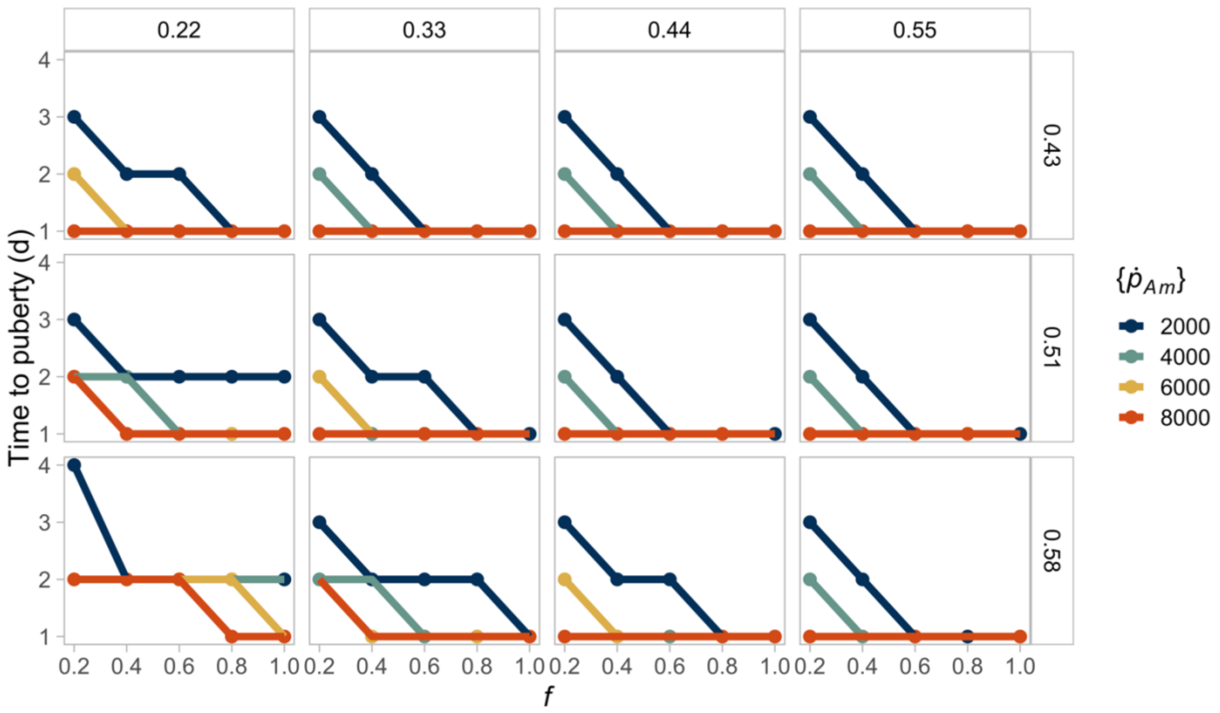


**Figure S4.** Interspecific differences in the time to reach puberty are amplified when a constant resource decreases. Individuals have faster developing times when the resource is non-limiting, and they combine high assimilation with high energy conductance. The columns indicate the value of energy conductance, while the rows represent the fraction of energy allocated to soma. Point and line colours indicate the maximum specific assimilation rate value. Points and lines of the same colour in each box (equivalent to a parameter combination) represent the same species at different food conditions.

The relative differences across species are nearly constant at any constant resource level for reserve energy (Fig. S5) and are uniform for structural volume (Fig. S6). The small relative differences show that the resource has mainly a scaling effect on both variables. The relative differences in developmental time are not constant but remain small and likely not significant (Fig. S7).

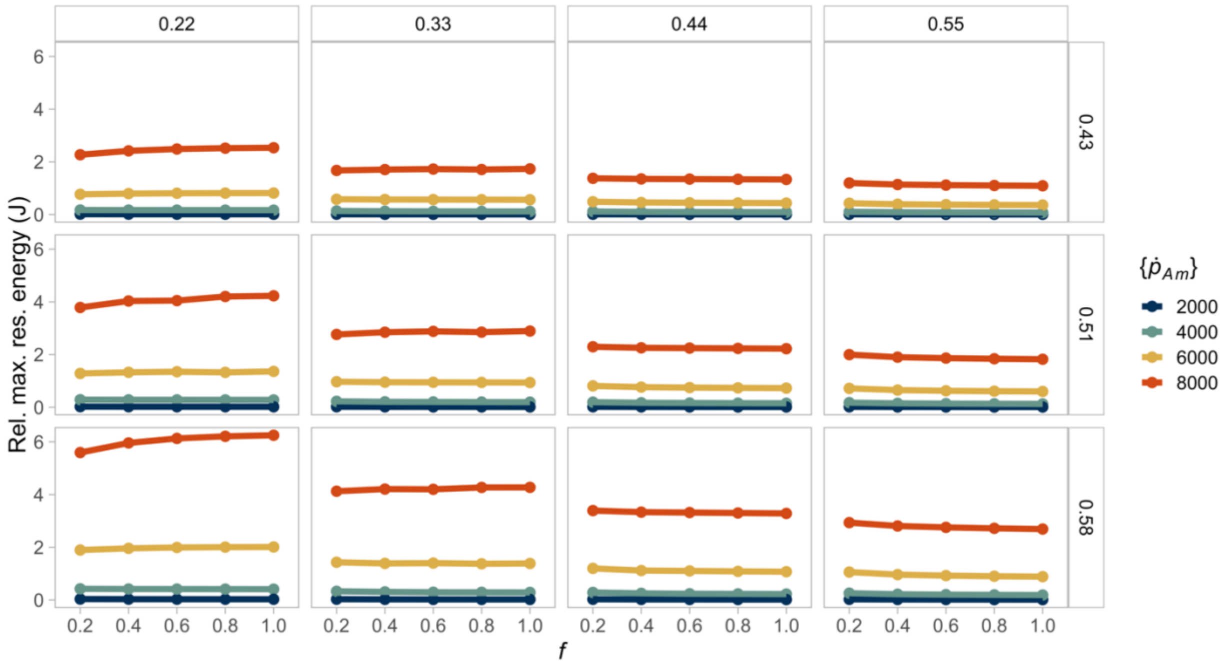


**Figure S5.** The resource scales the interspecific differences in reserve energy. Hence, differences between different constant food levels for the same species are small. The columns indicate the value of energy conductance, while the rows represent the fraction of energy allocated to soma. Point and line colours indicate the maximum specific assimilation rate value. Points and lines of the same colour in each box (equivalent to a parameter combination) represent the same species at different food conditions.

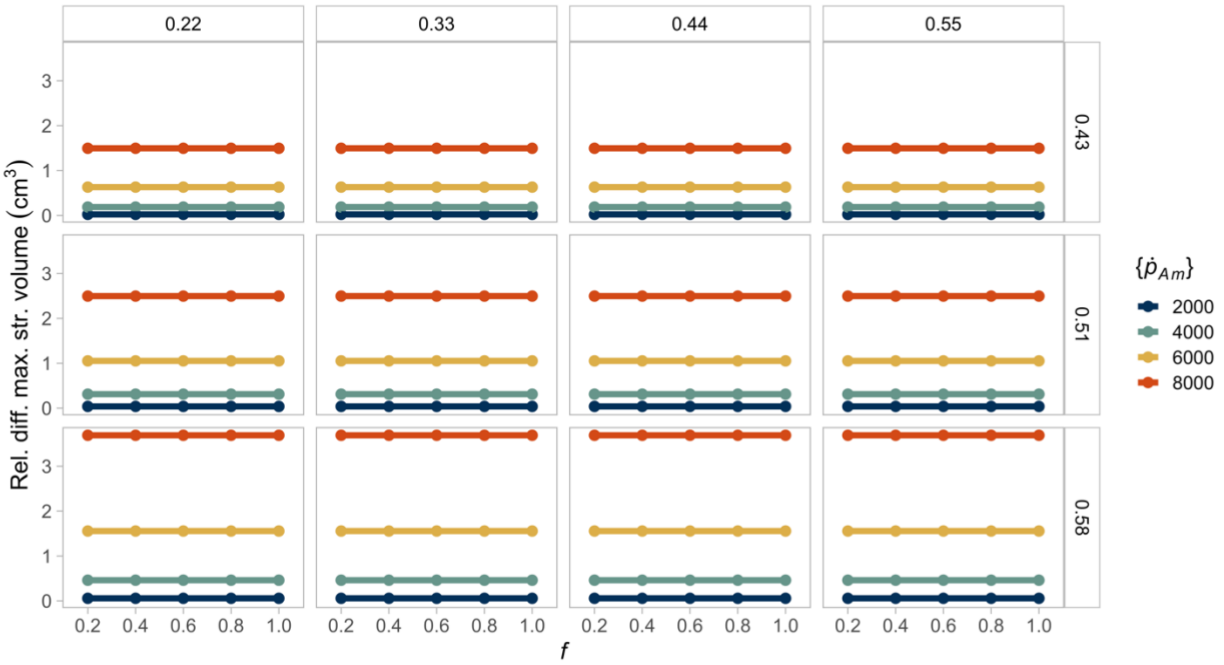


**Figure S6.** The resource scales the interspecific differences in structural volume. Hence, there are no relative differences between different constant food levels for the same species. The columns indicate the value of energy conductance, while the rows represent the fraction of energy allocated to soma. Point and line colours indicate the maximum specific assimilation rate value. Points and lines of the same colour in each box (equivalent to a parameter combination) represent the same species at different food conditions.

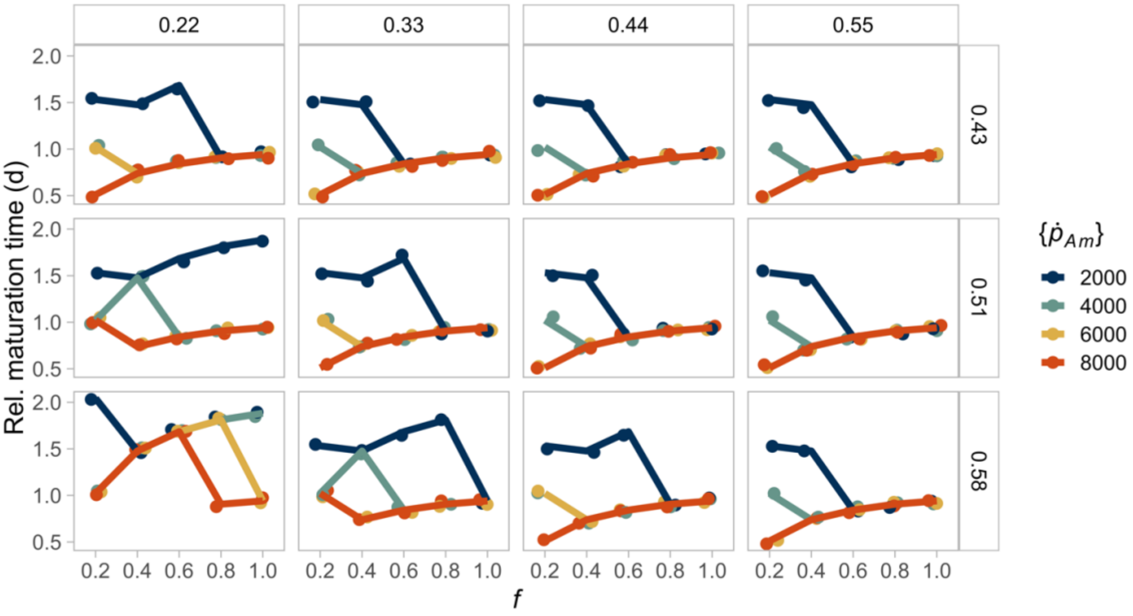


**Figure S7.** The relative differences between constant food levels in maturation time are small and likely not significant. The columns indicate the value of energy conductance, while the rows represent the fraction of energy allocated to soma. Point and line colours indicate the maximum specific assimilation rate value. Points and lines of the same colour in each box (equivalent to a parameter combination) represent the same species at different food conditions.

**Effect of resource variability**

A seasonal resource with a greater average amplifies the consequences of interspecific differences in biomass, reproductive output, reserve energy, and structural volume (Figs. S8 to S11, respectively). On the contrary, the interspecific differences in development time become greater as the mean resource decreases (Fig. S12). These differences in time to reach puberty are small and likely not significant. However, they depend on the initial food level or, equivalently, the birth time relative to the resource cycle.

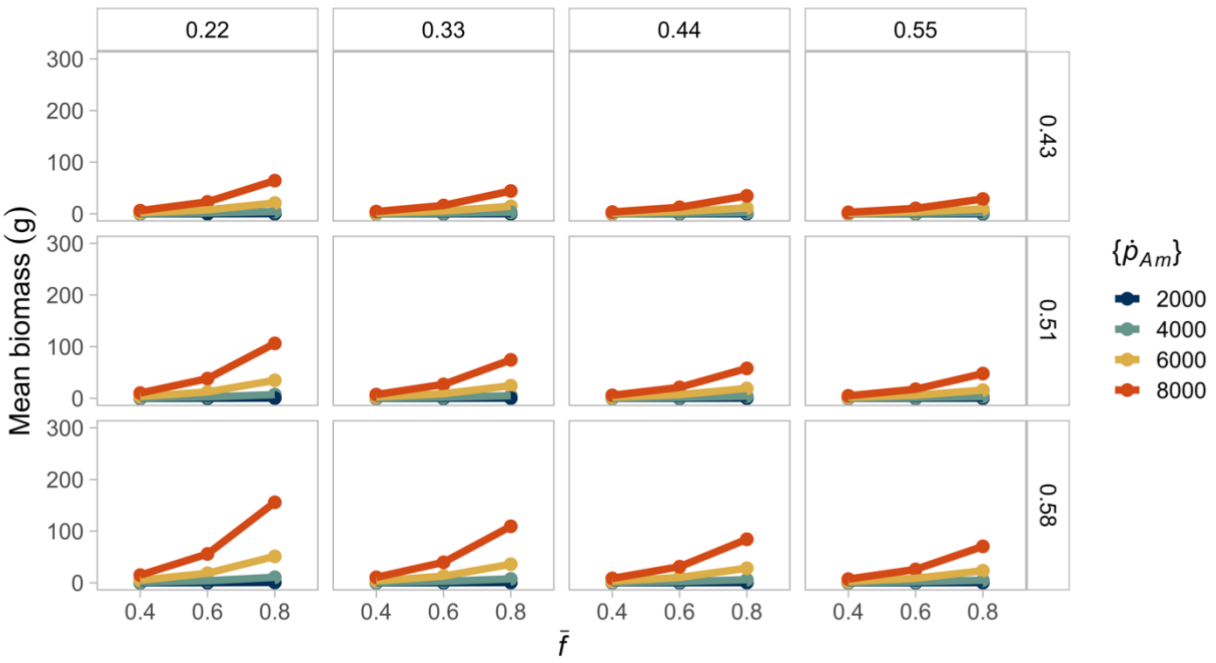


**Figure S8.** A seasonally varying resource with a lower average reduces interspecific differences in mean biomass, regardless of the initial food level. The largest biomass is attained when individuals combine high assimilation with low energy conductance. The columns indicate the value of energy conductance, while the rows represent the fraction of energy allocated to soma. Point and line colours indicate the maximum specific assimilation rate value. Points and lines of the same colour in each box (equivalent to a parameter combination) represent the same species at different food conditions. The value of $\bar{f}$ indicates the average resource level for each simulation. The initial resource level was set to the highest availability in each case.

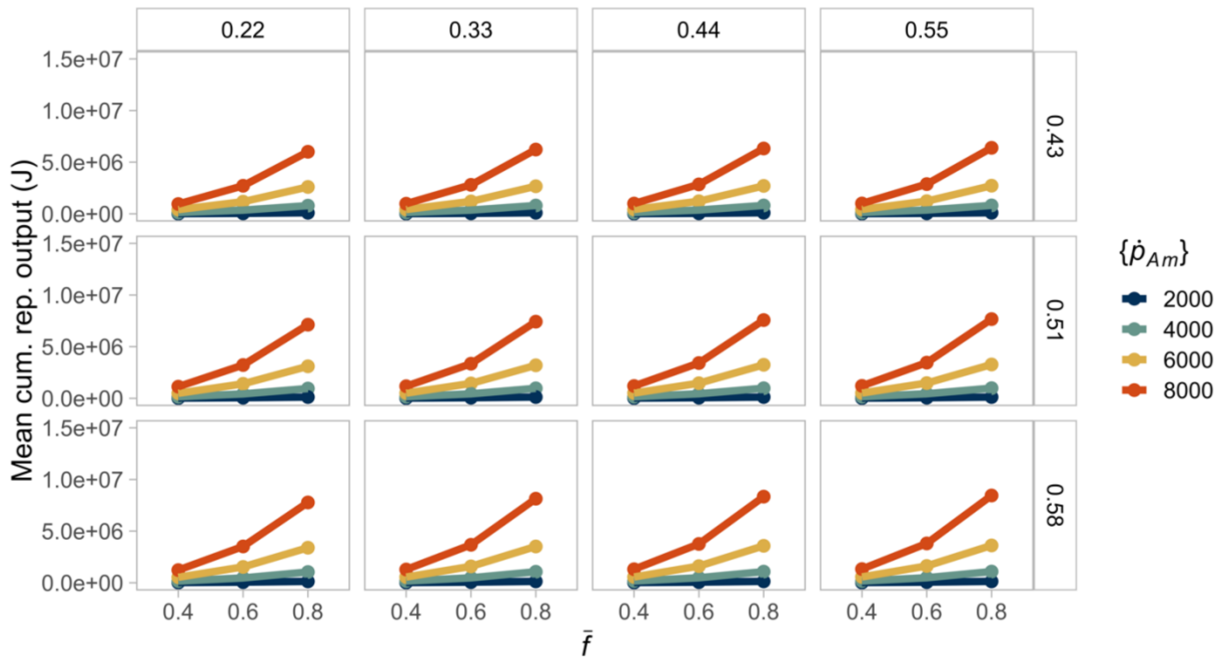


**Figure S9.** A seasonal and, on average scarcer resource reduces interspecific variability in mean cumulative reproductive output, regardless of the initial resource level. Higher reproductive output is reached when the fraction of energy allocated to soma is high. The columns indicate the value of energy conductance, while the rows represent the fraction of energy allocated to soma. Point and line colours indicate the maximum specific assimilation rate value. Points and lines of the same colour in each box (equivalent to a parameter combination) represent the same species at different food conditions. The value of $\bar{f}$ indicates the average resource level for each simulation. The initial resource level was set to the highest availability in each case (results are similar for all the initial food conditions, see figures S22 to S24).

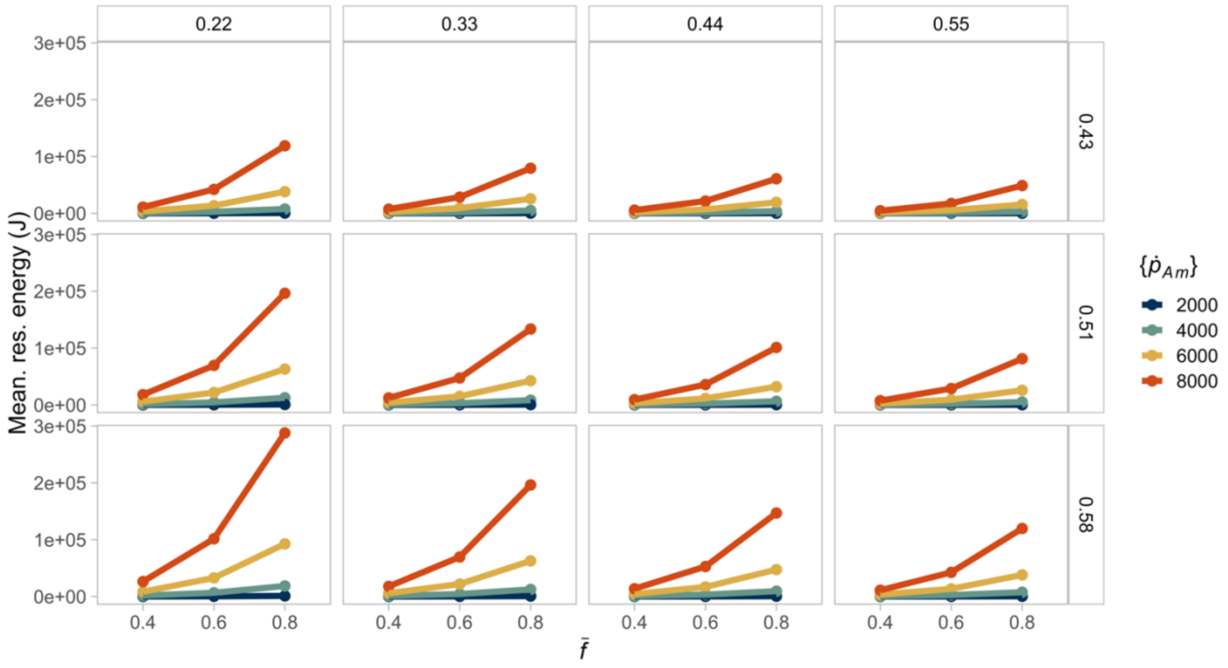


**Figure 10.** A seasonally varying resource with lower average reduces interspecific differences in mean reserve energy, regardless of the initial food level. The largest energy reserve is attained when individuals combine high assimilation with low energy conductance. The columns indicate the value of energy conductance, while the rows represent the fraction of energy allocated to soma. Point and line colours indicate the maximum specific assimilation rate value. Points and lines of the same colour in each box (equivalent to a parameter combination) represent the same species at different food conditions. The value of $\bar{f}$ indicates the average resource level for each simulation. The initial resource level was set to the highest availability in each case (results are similar for all the initial food conditions, see figures S13 to S15).

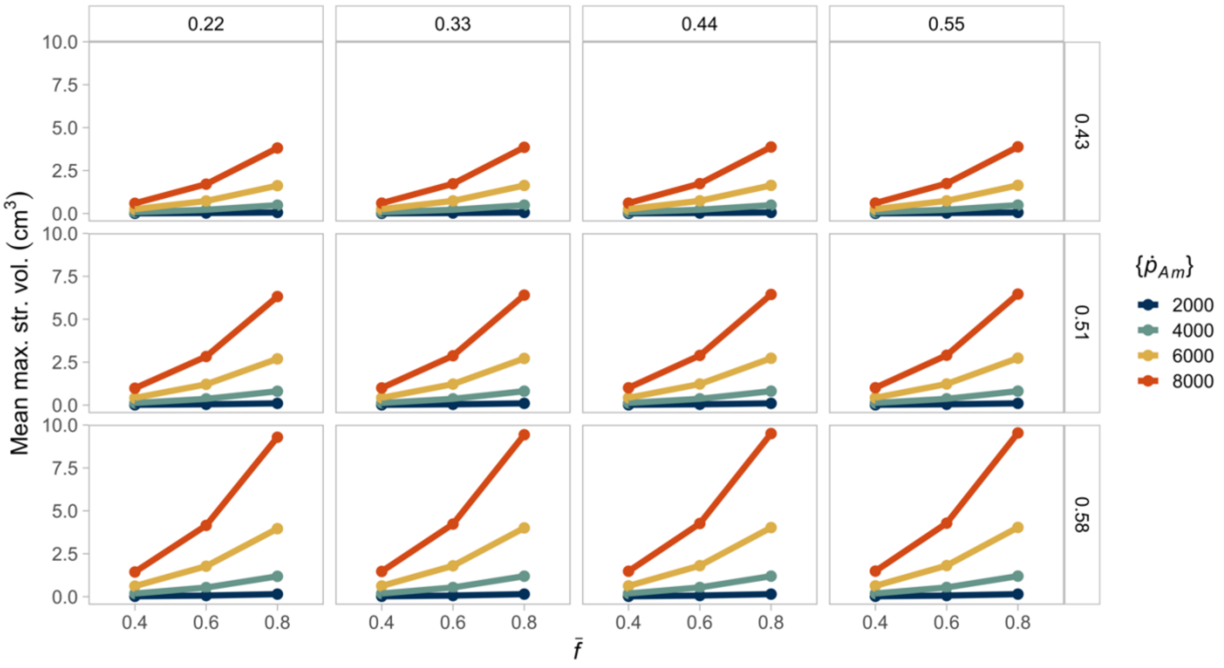


**Figure 11.** A seasonally varying resource with lower average reduces interspecific differences in mean structural volume, regardless of the initial food level. The largest volume is attained when individuals combine high assimilation with low energy conductance. The columns indicate the value of energy conductance, while the rows represent the fraction of energy allocated to soma. Point and line colours indicate the maximum specific assimilation rate value. Points and lines of the same colour in each box (equivalent to a parameter combination) represent the same species at different food conditions. The value of $\bar{f}$ indicates the average resource level for each simulation. The initial resource level was set to the highest availability in each case (results are similar for all the initial food conditions, see figures S16 to S18).

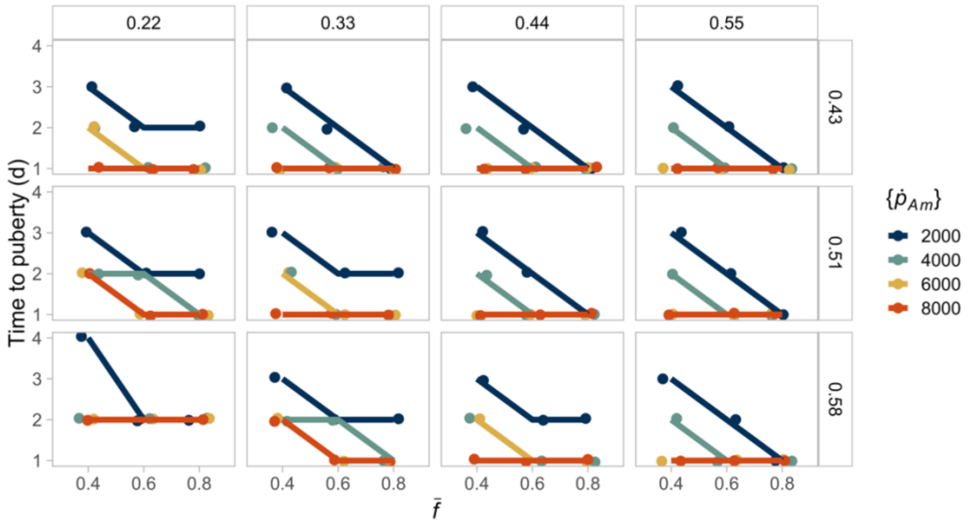


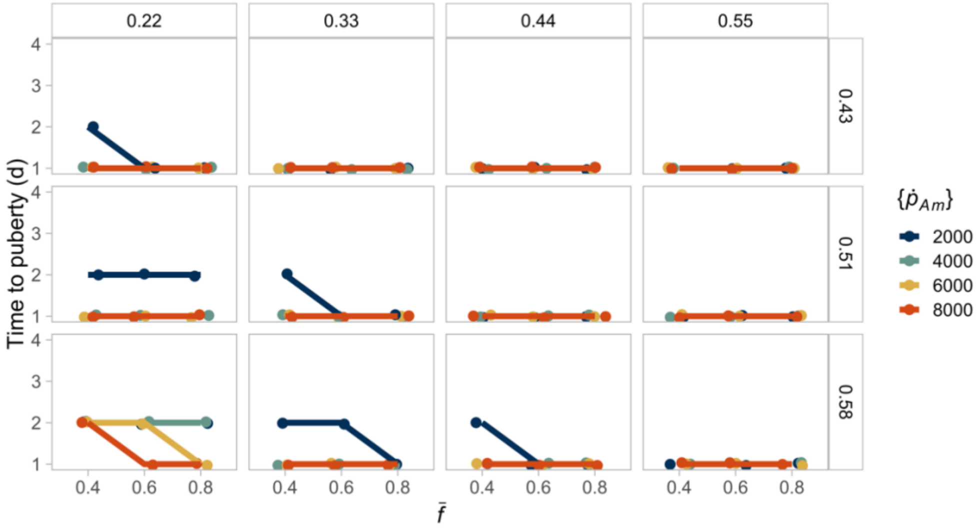


**Figure S12.** Interspecific differences in the time to reach puberty are amplified when the level of the resource is, on average, lower. A low initial resource produces individuals with slower development times (top) than the same individuals with maximum initial resource availability (bottom). In both scenarios, individuals develop faster when the resource is non-limiting, and they combine high assimilation with high energy conductance. The columns indicate the value of energy conductance, while the rows represent the fraction of energy allocated to soma. Point and line colours indicate the maximum specific assimilation rate value. Points and lines of the same colour in each box (equivalent to a parameter combination) represent the same species at different food conditions. The value of $\bar{f}$ indicates the average resource level for each simulation (see figures S22 to S24 for comparison among different initial resource conditions).

### Comparison across initial resource conditions

Regardless of the initial resource level, seasonality amplifies the consequences of interspecific differences in reserve energy (Figs. S13 to S15), structural volume (Figs. S16 to S18), and reproductive output (Figs. S19 to S21). The mean resource level seems to have a greater effect than the initial resource conditions on the development time (Figs. S22 to 24). However, the differences in time to reach puberty are likely not significant because their variation ranges from 2 to 4 days.

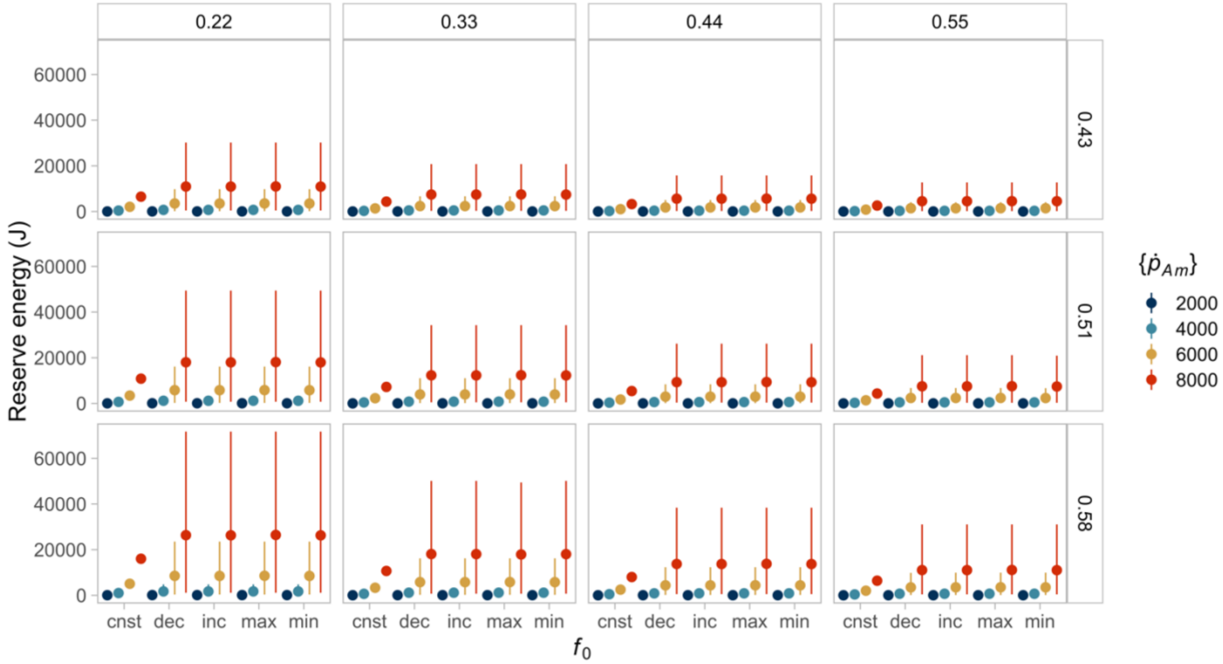


**Figure S13.** Individuals in a seasonal environment reach larger average reserve energy than the same individual in a constant environment with an equal mean resource availability, regardless of the initial resource condition $f_0$. Here, we compare individuals in a constant resource environment (cnst, $f = 0.4$) to those experiencing seasonality ($\bar{f} = 0.4$). We contrasted four different initial conditions for the seasonal environment, which means that individuals can be born when the resource is decreasing (dec, $f_0 = 0.3$), at the minimum level (min, $f_0 = 0.2$), increasing (inc, $f_0 = 0.3$) or at the maximum level (max, $f_0 = 0.4$). In the constant environment, points show the steady-state value reached at the end of the simulations (i.e., year three). In contrast, for the seasonal environment, points represent the average reserve energy calculated over the last two years (i.e., years one to three), and lines correspond to the minimum and maximum values. The columns indicate the value of energy conductance, while the rows represent the fraction of energy allocated to soma. Point and line colours indicate the maximum specific assimilation rate value. Points and lines of the same colour in each box (equivalent to a parameter combination) represent the same species at different food conditions.

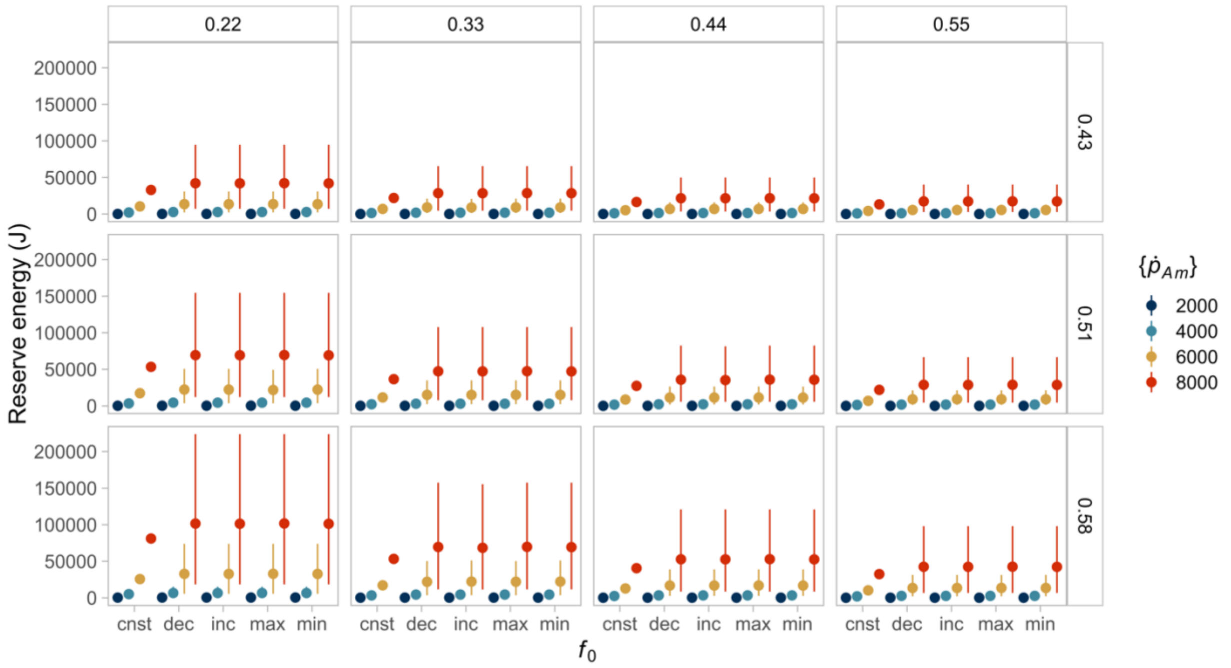


**Figure S14.** Individuals in a seasonal environment reach larger average reserve energy than the same individual in a constant environment with an equal mean resource availability, regardless of the initial resource condition $f_0$. Here, we compare individuals in a constant resource environment (cnst, $f = 0.6$) to those experiencing seasonality ($\bar{f} = 0.6$). We contrasted four different initial conditions for the seasonal environment, which means that individuals can be born when the resource is decreasing (dec, $f_0 = 0.5$), at the minimum level (min, $f_0 = 0.4$), increasing (inc, $f_0 = 0.5$) or at the maximum level (max, $f_0 = 0.8$). In the constant environment, points show the steady-state value reached at the end of the simulations (i.e., year three). In contrast, for the seasonal environment, points represent the average reserve energy calculated over the last two years (i.e., years one to three), and lines correspond to the minimum and maximum values. The columns indicate the value of energy conductance, while the rows represent the fraction of energy allocated to soma. Point and line colours indicate the maximum specific assimilation rate value. Points and lines of the same colour in each box (equivalent to a parameter combination) represent the same species at different food conditions.

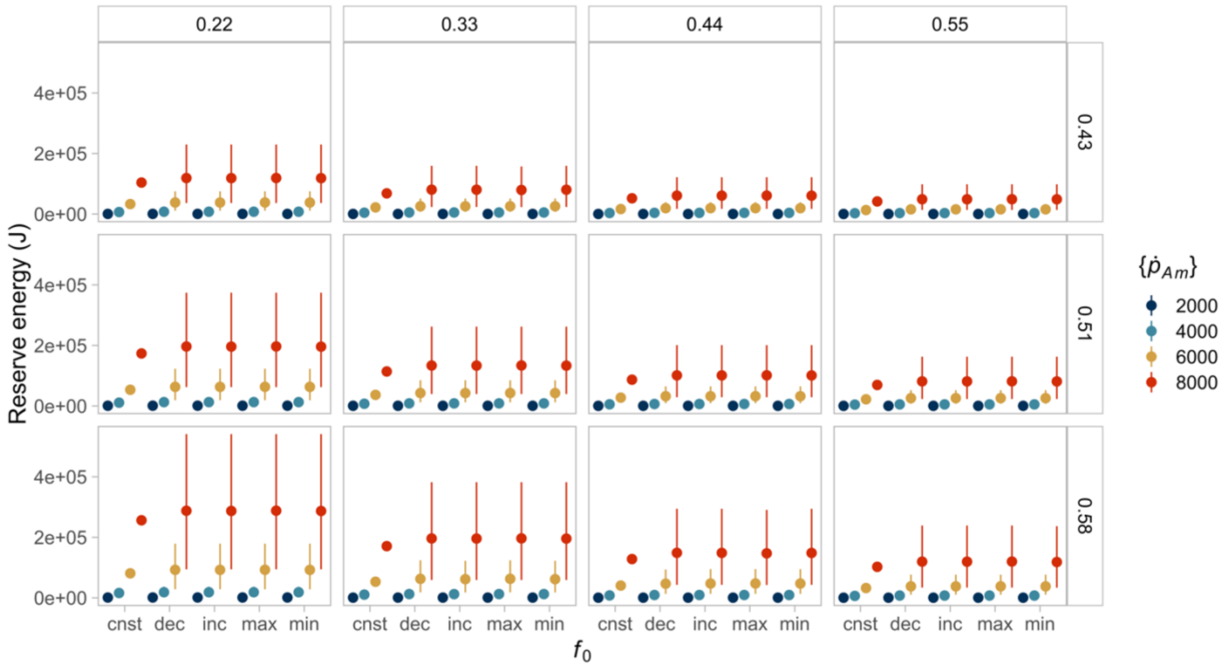


**Figure S15.** Individuals in a seasonal environment reach larger average reserve energy than the same individual in a constant environment with an equal mean resource availability, regardless of the initial resource condition $f_0$. Here, we compare individuals in a constant resource environment (cnst, $f = 0.8$) to those experiencing seasonality ($\bar{f} = 0.8$). We contrasted four different initial conditions for the seasonal environment, which means that individuals can be born when the resource is decreasing (dec, $f_0 = 0.7$), at the minimum level (min, $f_0 = 0.6$), increasing (inc, $f_0 = 0.7$) or at the maximum level (max, $f_0 = 1$). In the constant environment, points show the steady-state value reached at the end of the simulations (i.e., year three). In contrast, for the seasonal environment, points represent the average reserve energy calculated over the last two years (i.e., years one to three), and lines correspond to the minimum and maximum values. The columns indicate the value of energy conductance, while the rows represent the fraction of energy allocated to soma. Point and line colours indicate the maximum specific assimilation rate value. Points and lines of the same colour in each box (equivalent to a parameter combination) represent the same species at different food conditions.

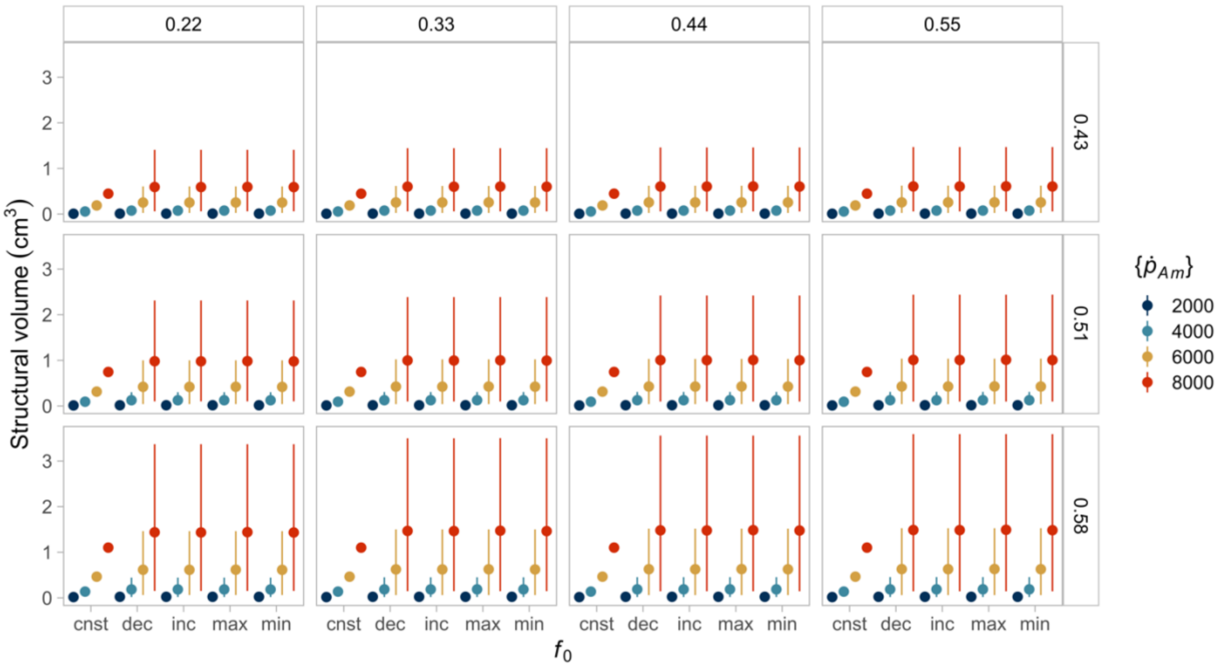


**Figure S16.** Individuals in a seasonal environment reach greater average structural volume than the same individual in a constant environment with an equal mean resource availability, regardless of the initial resource condition $f_0$. Here, we compare individuals in a constant resource environment (cnst, $f = 0.4$) to those experiencing seasonality ($\bar{f} = 0.4$). We contrasted four different initial conditions for the seasonal environment, which means that individuals can be born when the resource is decreasing (dec, $f_0 = 0.3$), at the minimum level (min, $f_0 = 0.2$), increasing (inc, $f_0 = 0.3$) or at the maximum level (max, $f_0 = 0.4$). In the constant environment, points show the steady-state value reached at the end of the simulations (i.e., year three). In contrast, for the seasonal environment, points represent the average structural volume calculated over the last two years (i.e., years one to three), and lines correspond to the minimum and maximum values. The columns indicate the value of energy conductance, while the rows represent the fraction of energy allocated to soma. Point and line colours indicate the maximum specific assimilation rate value. Points and lines of the same colour in each box (equivalent to a parameter combination) represent the same species at different food conditions.

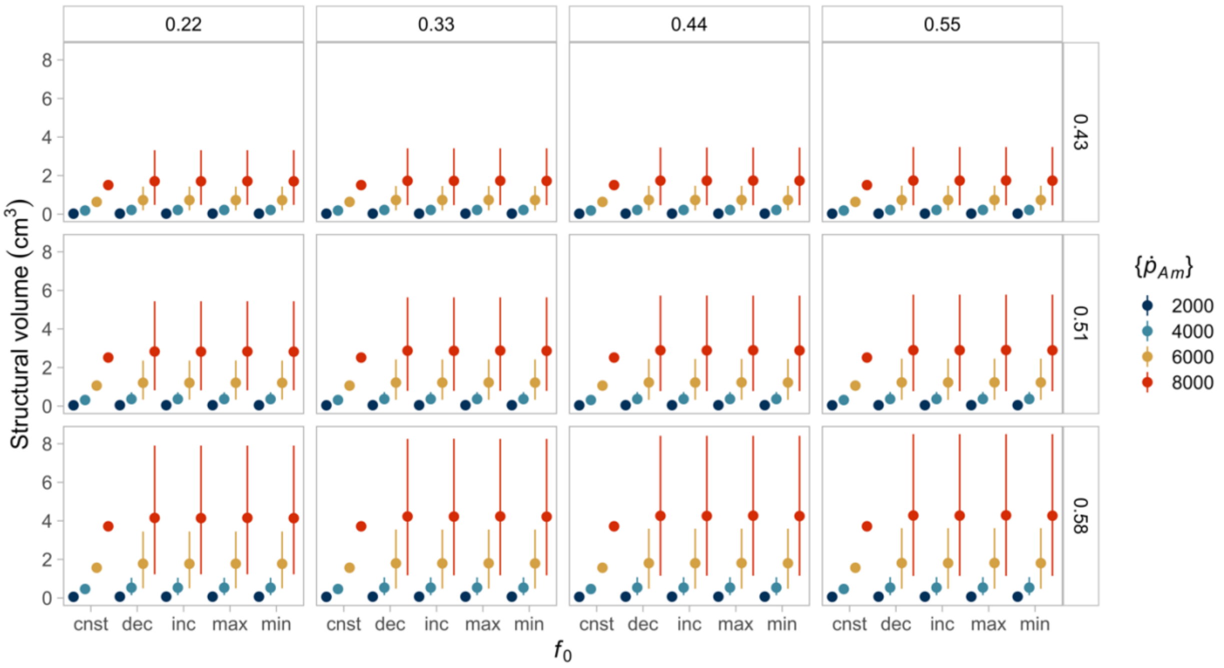


**Figure S17.** Individuals in a seasonal environment greater average structural volume than the same individual in a constant environment with an equal mean resource availability, regardless of the initial resource condition $f_0$. Here, we compare individuals in a constant resource environment (cnst, $f = 0.6$) to those experiencing seasonality ($\bar{f} = 0.6$). We contrasted four different initial conditions for the seasonal environment, which means that individuals can be born when the resource is decreasing (dec, $f_0 = 0.5$), at the minimum level (min, $f_0 = 0.4$), increasing (inc, $f_0 = 0.5$) or at the maximum level (max, $f_0 = 0.8$). In the constant environment, points show the steady-state value reached at the end of the simulations (i.e., year three). In contrast, for the seasonal environment, points represent the average structural volume calculated over the last two years (i.e., years one to three), and lines correspond to the minimum and maximum values. The columns indicate the value of energy conductance, while the rows represent the fraction of energy allocated to soma. Point and line colours indicate the maximum specific assimilation rate value. Points and lines of the same colour in each box (equivalent to a parameter combination) represent the same species at different food conditions.

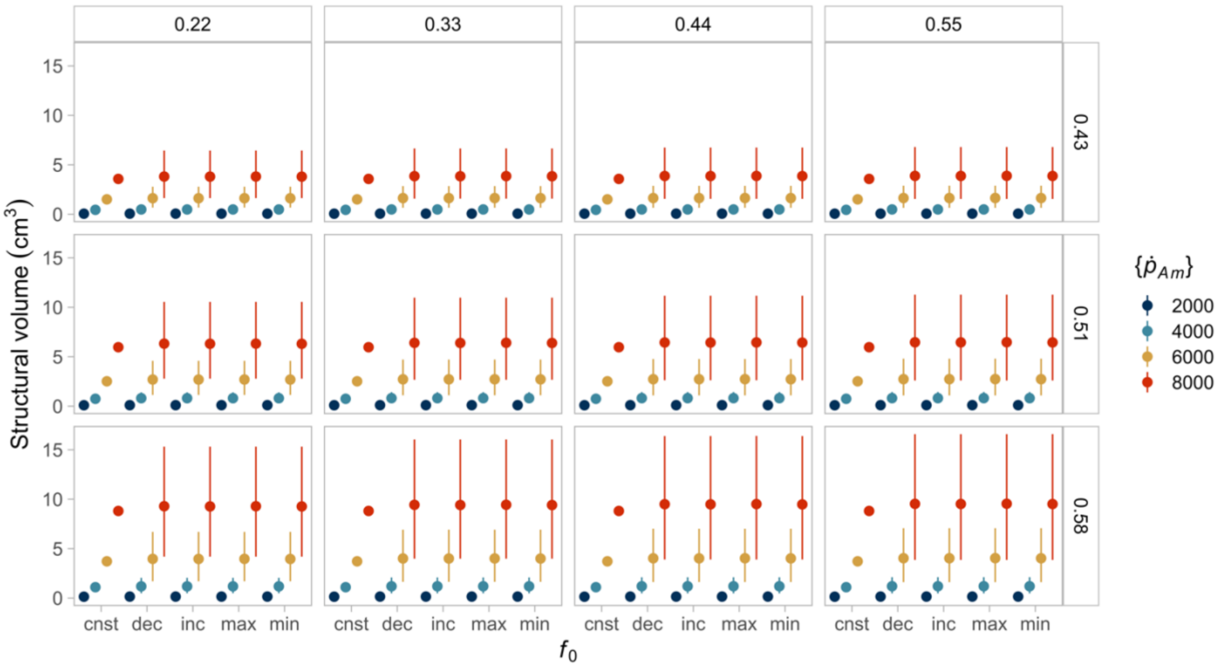


**Figure S18.** Individuals in a seasonal environment reach greater average structural volume than the same individual in a constant environment with an equal mean resource availability, regardless of the initial resource condition $f_0$. Here, we compare individuals in a constant resource environment (cnst, $f = 0.8$) to those experiencing seasonality ($\bar{f} = 0.8$). We contrasted four different initial conditions for the seasonal environment, which means that individuals can be born when the resource is decreasing (dec, $f_0 = 0.7$), at the minimum level (min, $f_0 = 0.6$), increasing (inc, $f_0 = 0.7$) or at the maximum level (max, $f_0 = 1$). In the constant environment, points show the steady-state value reached at the end of the simulations (i.e., year three). In contrast, for the seasonal environment, points represent the average structural volume calculated over the last two years (i.e., years one to three), and lines correspond to the minimum and maximum values. The columns indicate the value of energy conductance, while the rows represent the fraction of energy allocated to soma. Point and line colours indicate the maximum specific assimilation rate value. Points and lines of the same colour in each box (equivalent to a parameter combination) represent the same species at different food conditions.

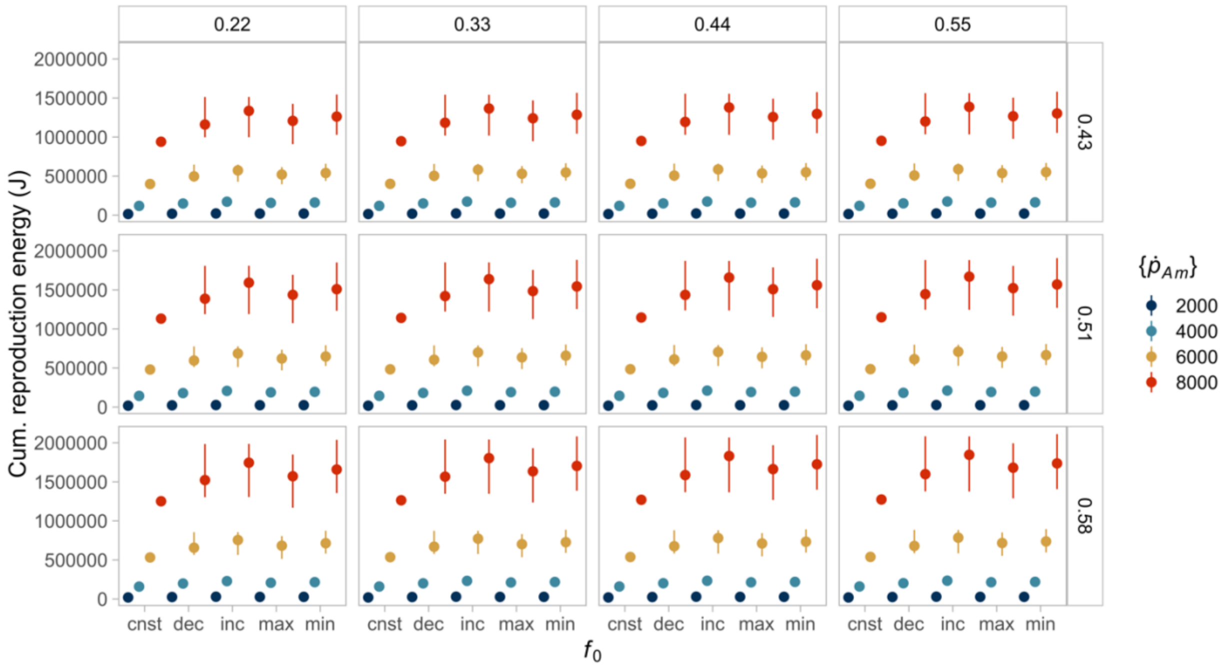


**Figure S19.** Individuals in a seasonal environment reach greater average cumulative reproduction energy than the same individual in a constant environment with an equal mean resource availability, regardless of the initial resource condition $f_0$. Here, we compare individuals in a constant resource environment (cnst, $f = 0.4$) to those experiencing seasonality ($\bar{f} = 0.4$). We contrasted four different initial conditions for the seasonal environment, which means that individuals can be born when the resource is decreasing (dec, $f_0 = 0.3$), at the minimum level (min, $f_0 = 0.2$), increasing (inc, $f_0 = 0.3$) or at the maximum level (max, $f_0 = 0.4$). In all environments, points show the average reproductive output calculated over the last two years (i.e., years one to three), and lines (for seasonal environments only) correspond to the minimum and maximum values. The columns indicate the value of energy conductance, while the rows represent the fraction of energy allocated to soma. Point and line colours indicate the maximum specific assimilation rate value. Points and lines of the same colour in each box (equivalent to a parameter combination) represent the same species at different food conditions.

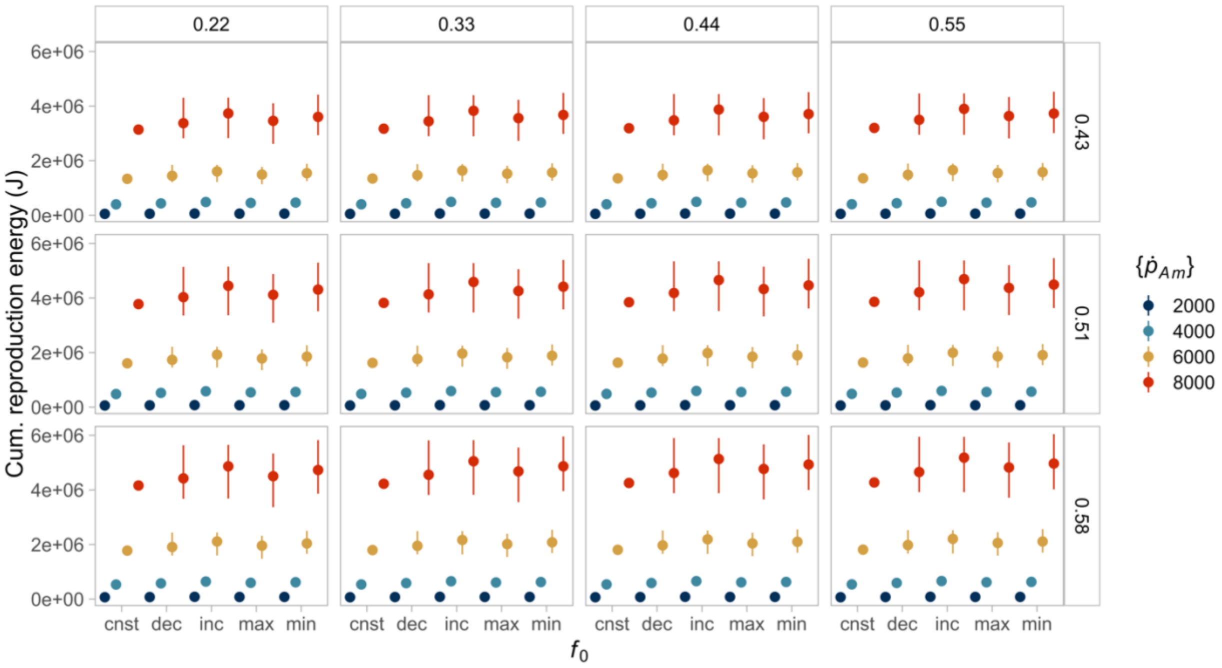


**Figure S20.** Individuals in a seasonal environment greater average cumulative reproduction energy than the same individual in a constant environment with an equal mean resource availability, regardless of the initial resource condition $f_0$. Here, we compare individuals in a constant resource environment (cnst, $f = 0.6$) to those experiencing seasonality ($\bar{f} = 0.6$). We contrasted four different initial conditions for the seasonal environment, which means that individuals can be born when the resource is decreasing (dec, $f_0 = 0.5$), at the minimum level (min, $f_0 = 0.4$), increasing (inc, $f_0 = 0.5$) or at the maximum level (max, $f_0 = 0.8$). In all environments, points show the average reproductive output calculated over the last two years (i.e., years one to three), and lines (for seasonal environments only) correspond to the minimum and maximum values. The columns indicate the value of energy conductance, while the rows represent the fraction of energy allocated to soma. Point and line colours indicate the maximum specific assimilation rate value. Points and lines of the same colour in each box (equivalent to a parameter combination) represent the same species at different food conditions.

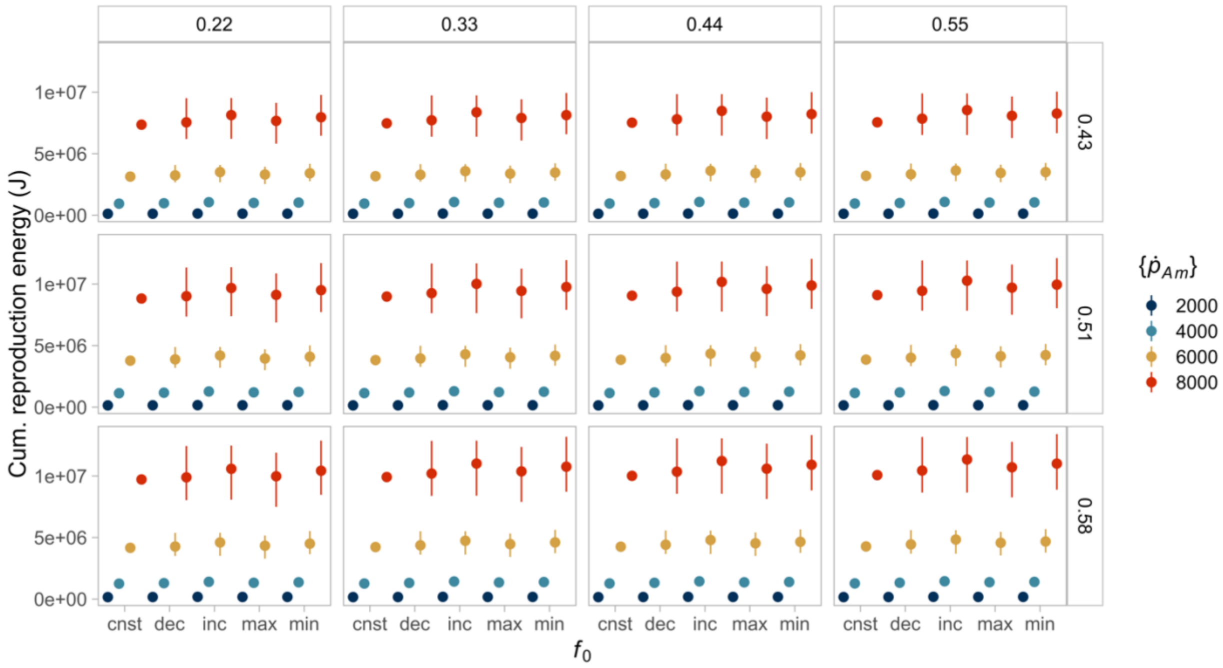


**Figure S21.** Individuals in a seasonal environment reach greater average cumulative reproduction energy than the same individual in a constant environment with an equal mean resource availability, regardless of the initial resource condition $f_0$. Here, we compare individuals in a constant resource environment (cnst, $f = 0.8$) to those experiencing seasonality ($\bar{f} = 0.8$). We contrasted four different initial conditions for the seasonal environment, which means that individuals can be born when the resource is decreasing (dec, $f_0 = 0.7$), at the minimum level (min, $f_0 = 0.6$), increasing (inc, $f_0 = 0.7$) or at the maximum level (max, $f_0 = 1$). In all environments, points show the average reproductive output calculated over the last two years (i.e., years one to three), and lines (for seasonal environments only) correspond to the minimum and maximum values. The columns indicate the value of energy conductance, while the rows represent the fraction of energy allocated to soma. Point and line colours indicate the maximum specific assimilation rate value. Points and lines of the same colour in each box (equivalent to a parameter combination) represent the same species at different food conditions.

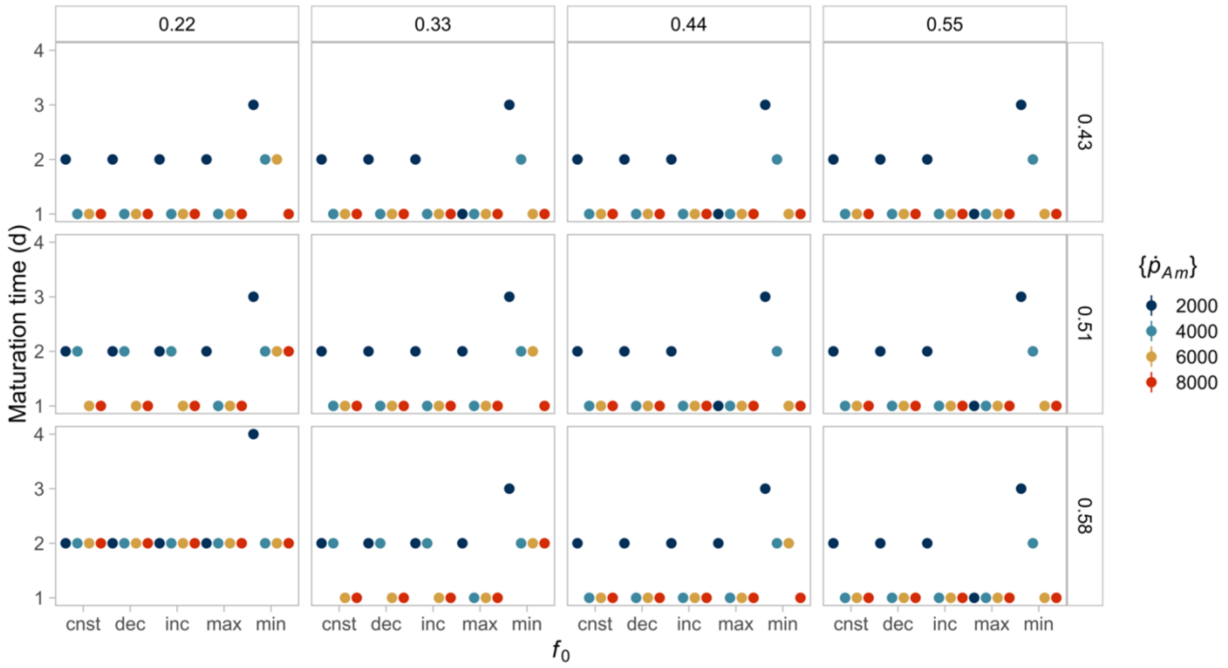


**Figure S22.** Interspecific differences in the time to reach puberty between individuals in a seasonal environment and the same individual in a constant environment with an equal mean resource availability do not seem to be dependent on the initial resource condition, $f_0$. Here, we compare individuals in a constant resource environment (cnst, $f = 0.4$) to those experiencing seasonality ($\bar{f} = 0.4$). We contrasted four different initial conditions for the seasonal environment, which means that individuals can be born when the resource is decreasing (dec, $f_0 = 0.3$), at the minimum level (min, $f_0 = 0.2$), increasing (inc, $f_0 = 0.3$) or at the maximum level (max, $f_0 = 0.6$). In all environments, points show the numbers of days required to reach the puberty threshold ($E_H^p$). The columns indicate the value of energy conductance, while the rows represent the fraction of energy allocated to soma. Point colours indicate the maximum specific assimilation rate value. Points of the same colour in each box (equivalent to a parameter combination) represent the same species at different food conditions.

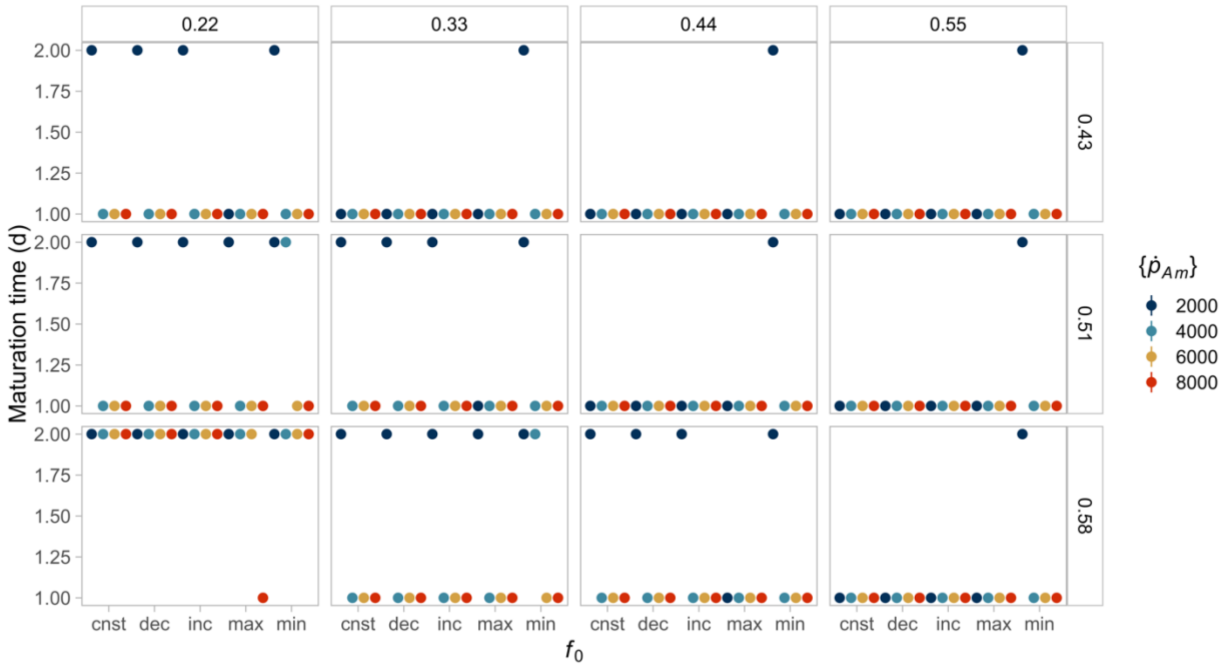


**Figure S23.** Interspecific differences in the time to reach puberty between individuals in a seasonal environment and the same individual in a constant environment with an equal mean resource availability do not seem to be dependent on the initial resource condition, $f_0$. Here, we compare individuals in a constant resource environment (cnst, $f = 0.6$) to those experiencing seasonality ($\bar{f} = 0.6$). We contrasted four different initial conditions for the seasonal environment, which means that individuals can be born when the resource is decreasing (dec, $f_0 = 0.5$), at the minimum level (min, $f_0 = 0.4$), increasing (inc, $f_0 = 0.5$) or at the maximum level (max, $f_0 = 0.8$). In all environments, points show the numbers of days required to reach the puberty threshold ($E_H^p$). The columns indicate the value of energy conductance, while the rows represent the fraction of energy allocated to soma. Point colours indicate the maximum specific assimilation rate value. Points of the same colour in each box (equivalent to a parameter combination) represent the same species at different food conditions.

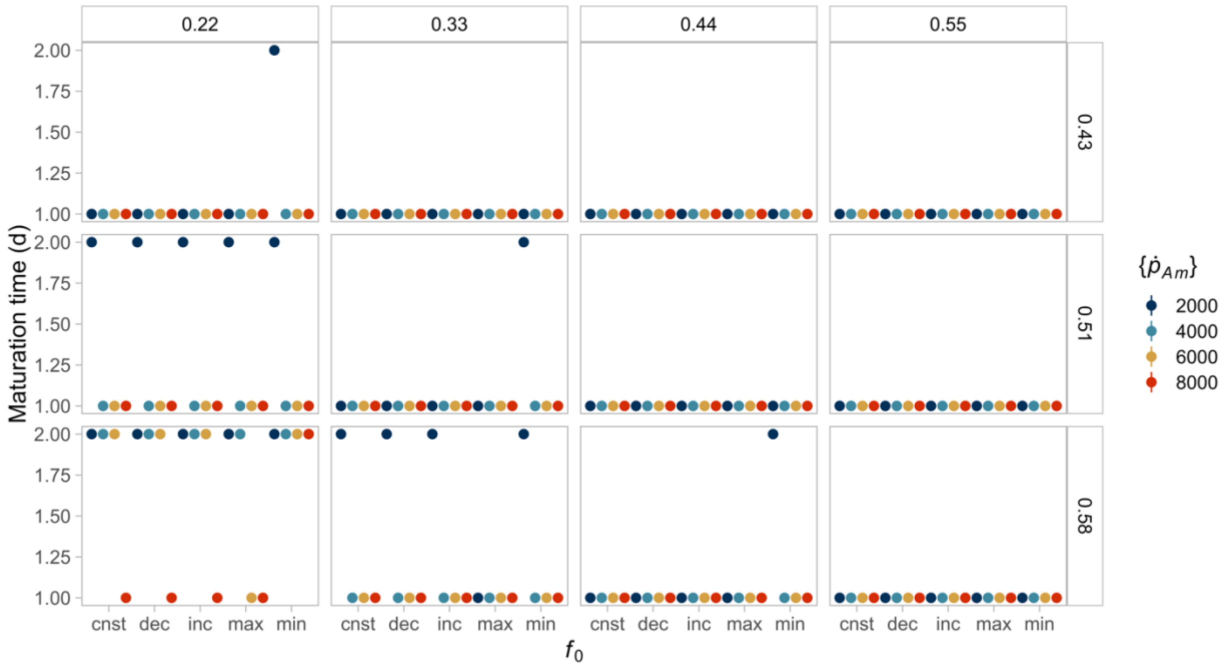


**Figure S24.** Interspecific differences in the time to reach puberty between individuals in a seasonal environment and the same individual in a constant environment with an equal mean resource availability do not seem to be dependent on the initial resource condition, $f_0$. Here, we compare individuals in a constant resource environment (cnst, $f = 0.8$) to those experiencing seasonality ($\bar{f} = 0.8$). We contrasted four different initial conditions for the seasonal environment, which means that individuals can be born when the resource is decreasing (dec, $f_0 = 0.7$), at the minimum level (min, $f_0 = 0.6$), increasing (inc, $f_0 = 0.7$) or at the maximum level (max, $f_0 = 1$). In all environments, points show the numbers of days required to reach the puberty threshold ($E_H^p$). The columns indicate the value of energy conductance, while the rows represent the fraction of energy allocated to soma. Point colours indicate the maximum specific assimilation rate value. Points of the same colour in each box (equivalent to a parameter combination) represent the same species at different food conditions.

### Comparing temperate and tropical species: an example

We used the Fan-tailed Gerygone, *Gerygone flavolateralis,* and the Grey Warbler, *G. igata*, to compare the traits exhibited by related species in contrasting environments. We chose these species because they were originally within our parameter space. However, we included more data and reestimated their parameters to improve the accuracy of the predictions (Tab. S2).

**Table S2.** Parameter values for the two species used to compare tropical versus temperate species.

| Species | Common name | $\kappa$ | $\{\dot{p}_{Am}\}$ | $\dot{v}$ | $[\dot{p}_M]$ | $[E_G]$ | $E_H^b$ | $E_H^p$ |
|---|---|---|---|---|---|---|---|---|
| *Gerygone flavolateralis* | Fan-tailed Gerygone | 0.998 | 4692.6 | 0.08 | 5084.7 | 7338 | 7.647 | 453.1 |
| *Gerygone igata* | Grey Warbler | 0.983 | 4878.1 | 0.05 | 6045.1 | 7351 | 71.74 | 7880 |
